# *In silico* identification of potential inhibitors against human 2’-5’-oligoadenylate synthetase (OAS) proteins

**DOI:** 10.1101/804716

**Authors:** Karen J. Gonzalez, Diego Moncada-Giraldo, Juan B. Gutierrez

## Abstract

As part of the type I IFN signaling, the 2’-5’- oligoadenylate synthetase (OAS) proteins have been involved in the progression of several non-viral diseases. Notably, OAS has been correlated with immune-modulatory functions that promote chronic inflammatory conditions, autoimmune disorders, cancer, and infectious diseases. In spite of this, OAS enzymes have been ignored as drug targets, and to date, there are no reports of compounds that can inhibit their activity. In this study, we have used homology modeling and virtual high-throughput screening to identify potential inhibitors of the human proteins OAS1, OAS2, and OAS3. Altogether, we have found 37 molecules that could exert a competitive inhibition in the ATP binding sites of OAS proteins, independently of the activation state of the enzyme. This latter characteristic, which might be crucial for a versatile inhibitor, was observed in compounds interacting with the residues Asp75, Asp77, Gln229, and Tyr230 in OAS1, and their equivalents in OAS2 and OAS3. Although there was little correlation between specific chemical fragments and particular interactions, intermolecular contacts with OAS catalytic triad and other critical amino acids were mainly promoted by heterocycles with π electrons and hydrogen bond acceptors. In conclusion, this study provides a potential set of OAS inhibitors as well as valuable information for their design, development, and optimization.

## Introduction

Inflammation is the defense mechanism employed by the immune system to counter potentially harmful agents [1]. During viral infections, inflammatory responses are initiated by the activation of specific molecular sensors—known as pattern-recognition receptors (PRRs)—that detect pathogen-associated molecular patterns (PAMPs) [2]. Unlike bacteria, where PAMPs constitute a broad set of molecules, viral PAMPs consist mainly of nucleic acids and viral fusion glycoproteins [3]. As a result of PRRs activity, type I interferons (IFN) and other mediators of the innate immune response are produced to exert an antiviral function [4].

In the presence of double-stranded (ds) RNA viruses, interferon-stimulated genes such as the 2’-5’-oligoadenylate synthetase (OAS) are activated to detect and restrict viral replication [5], [6]. In humans, the OAS family consists of three catalytically active enzymes, OAS1, OAS2, OAS3, and one OAS-like protein lacking oligoadenylate synthase activity [7]. Whereas OASL exerts its antiviral role through a ubiquitin-like domain [8], the remaining family members use a nucleotidyltransferase domain to catalyze the conversion of ATP into 2’-5’ -linked oligoadenylates (2-5A) [9]. These 2-5A act as unique second messengers that trigger the activation of the ribonuclease L and induce the degradation of cellular and viral RNA [10].

The activation mechanism of OAS has been shown to involve the recognition of dsRNA, followed by a significant structural rearrangement on the protein [11]-[13]. Specifically, the inactive conformation of the enzyme senses dsRNA through a channel of positively charged residues located opposite to the active site [12], [13]. As a result of this interaction, OAS experiences a conformational change in its catalytic domain, bringing the active site into proximity and acquiring an optimal configuration for the catalytic reaction [13]. Although the mechanistic details of OAS activation have been studied mainly in OAS1, it is believed that the process is conserved for both OAS2 and OAS3 [6].

After stimulation with dsRNA, OAS requires two Mg^2+^ ions and two molecules of ATP to perform its enzymatic activity [14]. During the synthesis of 2-5A, one ATP, known as the donor, transfers its AMP moiety to a second ATP, known as the acceptor. For polymerization, the acceptor ATP is replaced by a growing chain of 2-5A [15]. Since the 2’-specificity of the oligomer is controlled by the binding mode of the acceptor ATP [12], the Mg^2+^ ions play a critical role in positioning the substrates in an ideal geometry for the reaction. Furthermore, Mg^2+^ is also responsible for increasing the reactivity of the 2’ oxygen of the acceptor ATP [12]. The binding of ATP and two Mg^2+^ ions leave the OAS proteins fully activated with a structure referred to as the active conformation [12].

Although the OAS family of proteins is known as an essential player in the effector response against viral infections [16]. OAS has also been correlated with non-viral diseases such as chronic inflammatory conditions [17], [18], autoimmune disorders [19]–[21], oral cancer [22], tuberculosis [23], and malaria [24]. As a common characteristic, OAS has been described as part of the type I IFN signaling, exerting a significant immunoregulatory role. Besides being involved in the progression of several diseases, OAS proteins have also been shown to contribute with resistance to the only approved treatment for gastric cancer [25] and oncolytic virus therapies for pancreatic ductal adenocarcinoma (PDA) [26].

In oral cancer, it has been demonstrated that high expression of OAS2 promotes tumor progression by negative modulation of the T-cell functions. Particularly, OAS2 can regulate the TCR-ζ chain (CD3 ζ) expression and with it, limit the response required for controlling tumor growth [22]. Other diseases associated with CD3-ζ chain expression have also shown up-regulation of OAS genes. In systemic lupus erythematosus (SLE), an autoimmune disorder related to the deficiency of CD3-ζ chain expression, increased levels of OAS were found to be correlated with disease activity [21], [27]. Likewise, although no association has been confirmed with clinical parameters, the up-regulation of OAS was observed in relapsing-remitting multiple sclerosis [28] and systemic sclerosis [29].

In pancreatic β cells, high activation of OAS proteins increases basal apoptosis and augment the risk of developing type 1 diabetes (T1D) [19]. This relationship was also found at the genotype level, where individuals with T1D had higher frequencies of polymorphisms associated with abnormal enzymatic activity than nondiabetic subjects [20]. Regarding infectious diseases, OAS expression has been reported as a signature that differentiates active tuberculosis from latent tuberculosis. As a consequence, it has been proposed that OAS has an immune-modulatory function that contributes to disease progression [23]. Similarly, the analysis of the OAS role during malaria infections has demonstrated that deficient suppression of type I IFN signaling may promote severe infections [24].

The critical role of OAS in non-viral diseases shows that this family of proteins might be a new target for immune modulation [18] or improvement of existing treatments. Nonetheless, this approach has gone unnoticed in the drug discovery field, and to date, there are no reports of small compounds that can prevent OAS activity. Consequently, this study aims at the *in-silico* identification of potential OAS inhibitors. By virtual high-throughput screening, we identified 37 small drug-like molecules that could block all substrate binding sites of OAS and limit its 2-5A synthesis activity.

## Methodology

### Structural modeling

The nucleotidyltransferase domains of the human proteins OAS1, OAS2, and OAS3 were used as targets for virtual screening. The crystal structure of OAS1 was retrieved from the Protein Data Bank [30], [31], with the PDB identifier <u>4IG8</u> [13]. In the absence of crystal structures for OAS2 and OAS3, these proteins were modeled by homology modeling. The sequences of OAS2 and OAS3 were obtained from NCBI [32] (accession codes <u>NP 058197.2,</u> and <u>NP 006178.2</u>), and their domains were defined with the UniProt database [33] (UniProt ID: <u>P29728,</u> and <u>Q9Y6K5</u>). Using l-TASSER [34]–[36], the 3D structures of OAS2 and OAS3 were constructed considering the usual active and inactive conformations of OAS proteins. The template used for modeling was the porcine OAS1 (pOAS1) in its active and inactive (PDB:) states (PDB: <u>4RWN</u> and <u>1PX5</u>) [12]. Since the crystal structure of human OAS1 is only available in an active conformation, the inactive state of this enzyme was also modeled. The template pOAS1 has 75.1, 53.1, and 56.6% sequence identity with human OAS1, OAS2, and OAS3, respectively.

To consider the different activation states of OAS proteins during the virtual screening, the structures in an active conformation were turned into three models: active protein without ligands, active protein bound to Mg2+ ions, and active protein in complex with Mg2+ ions and a donor substrate. The holoenzymes were created according to pOAS1 (PDB: <u>4RWN</u>) as this enzyme has been crystallized with donor and acceptor ligands [12]. The donor substrate in the models corresponded to an ATP analog called α, β-Methyleneadenosine-5’-triphosphate (ApCpp). Altogether, twelve targets consisting of nine models and one crystal structure (counting for three activation states), were used for virtual screening (Table 1). An example of the 3D structures representing each activation state studied is presented in Figure 1.

**Figure 1.**
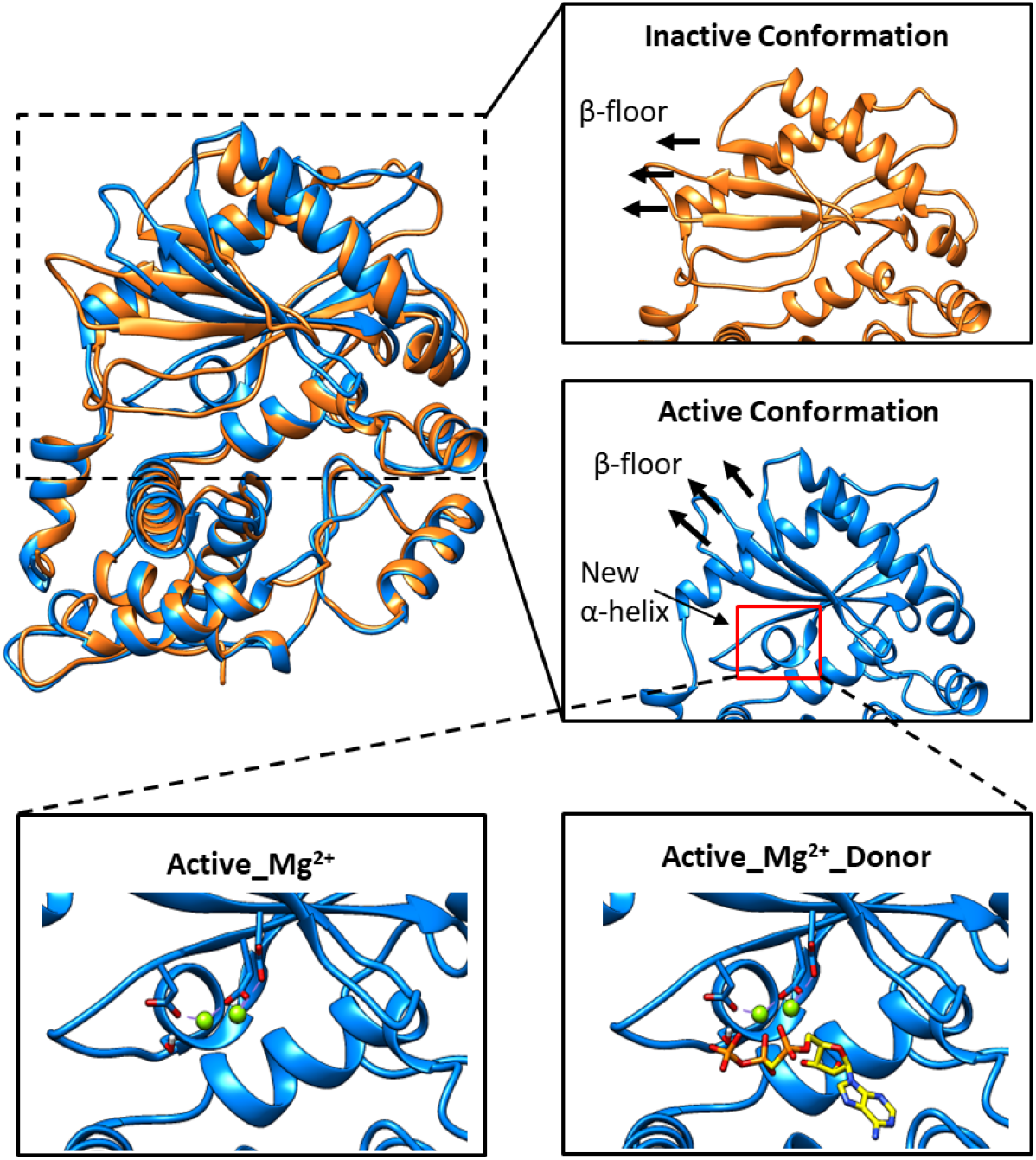
3D models representing the OAS activation states considered for virtual screening. On the top left is shown the structural alignment of the active (blue) and inactive (orange) conformations of human 0AS3. On the top right are displayed the conformational changes produced by the binding of RNA such as the movement of the β-floor (black arrows), and the formation of a new α-helix in the active conformation of the protein (red sguare) [11]—[13], [49]. To consider the catalytic mechanisms of OAS proteins, the active conformation is divided into three activation states representing the protein without ligands, the protein bound to Mg^2+^ ions, and the complex formed by Mg^2+^ ions and a donor substrate (ApCpp). Mg^2+^ ions are shown in green, and the ApCpp substrate is shown in yellow.

**Table 1.**
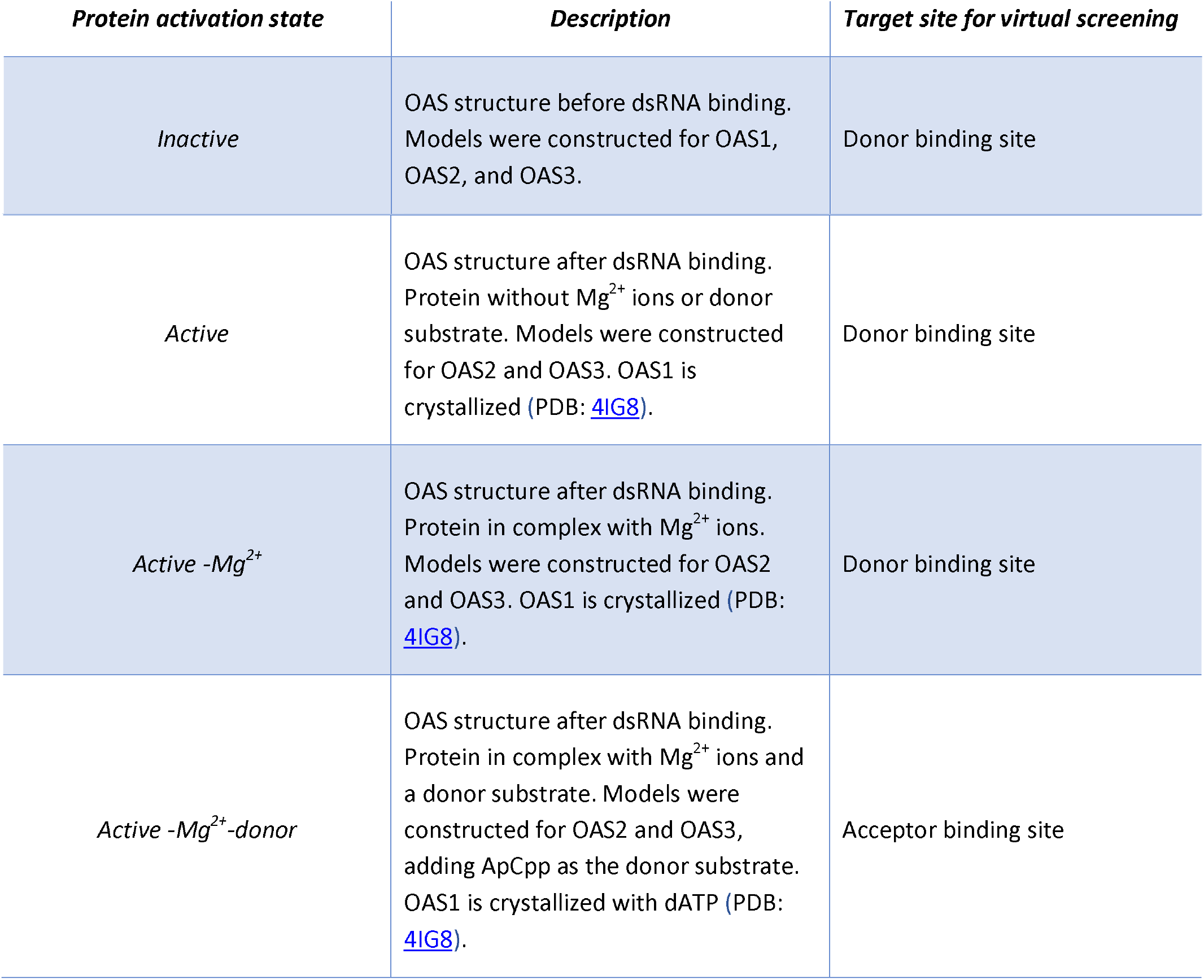
OAS activation states considered for virtual screening. Human proteins OAS1, OAS2, and OAS3 were studied in four different activation states derived from two structural conformations: Inactive, before dsRNA binding; and active, after dsRNA binding.

### *In silico* high-throughput screening

#### Target preparation for molecular docking

The crystal structure of human OAS1 <u>(4IG8</u>) was initially treated by removing crystallographic waters, cofactors, and ligands. Subsequently, all targets were prepared with the AutoDockTools 1-5.6 software [37], adding Gasteiger’s charges and polar hydrogens, specifying AutoDock atom types, and converting PDB files to PDBQT file formats.

#### Search space selection for virtual screening

Prior to the virtual screening, docking calculations with ATP and ATP analogs were performed on OAS1, OAS2, and OAS3 to identify search spaces that favored the natural binding mode of OAS substrates. Using the AutoDock Vina 1.1.2 software [38], different search volumes were evaluated around amino acids involved in OAS activity (Table 2) [12], [13], [39]. Important residues for OAS2 and OAS3 were established by sequence and structural alignments against human and porcine OAS1, using CLUSTAL O(1.2.4) [40] and UCSF Chimera 1.10.2 [41]. Docking calculations were carried out on all activation states of the proteins utilizing ATP, dATP, and ApCpp as substrates. The ATP structure was downloaded from the ZINC database [42], and dATP and ApCpp molecules were retrieved from the PDB files <u>4IG8</u> and <u>4RWN,</u> respectively. All the ligands were converted to the PDBQT format using the “prepare ligand” module available in AutoDockTools. The search spaces selected for virtual screening corresponded to search volumes where the substrates mimicked the binding modes reported in <u>4IG8</u> and <u>4RWN</u> (Figure S1). For the activation states that have not been crystallized with a ligand, the search spaces were chosen based on interactions with amino acids involved in OAS activity. The docking results with the native substrates were later used as positive controls to determine whether a molecule could be a competitive inhibitor (see Selection criteria section).

**Table 2.**
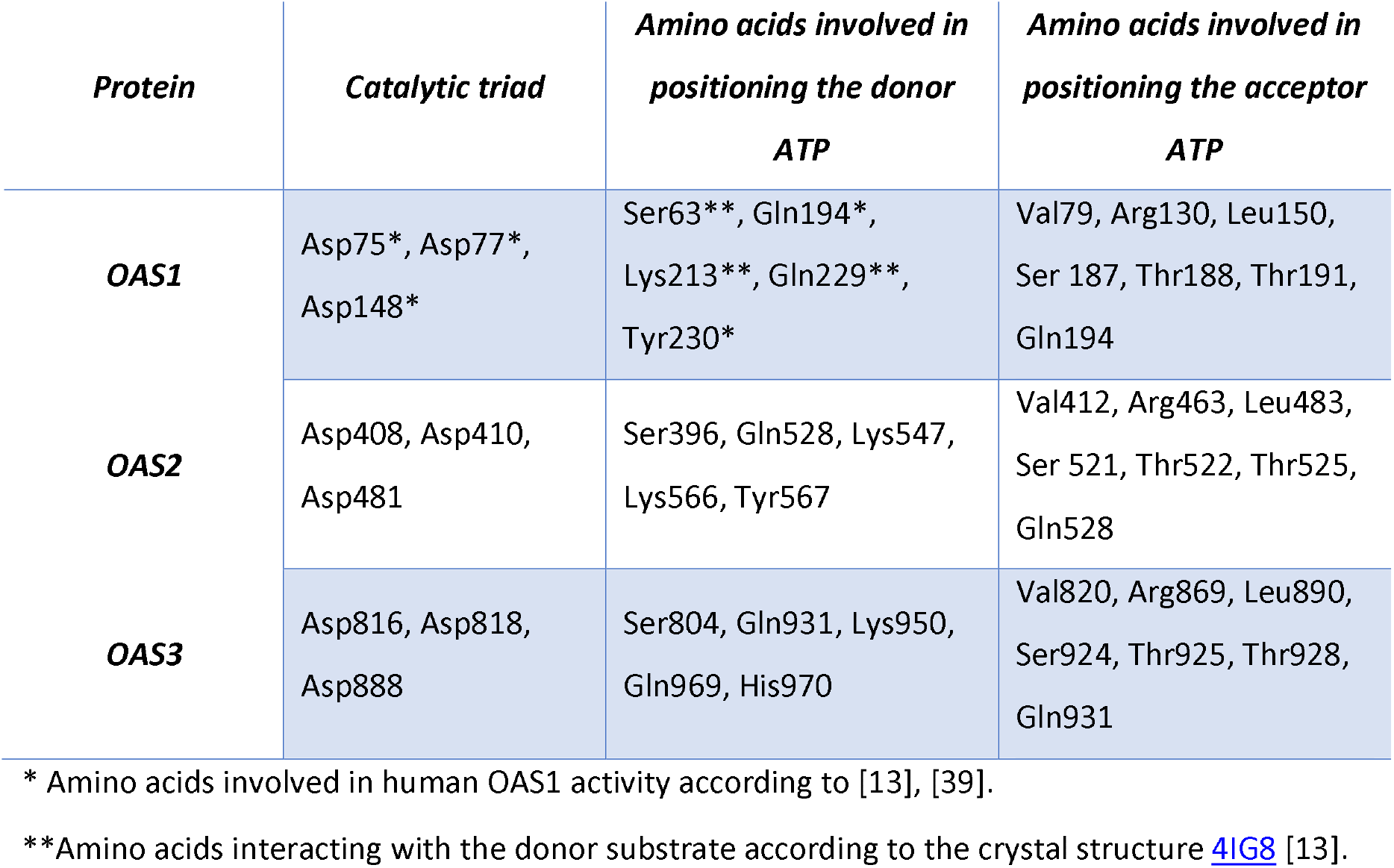
Amino acids involved in the enzymatic activity of human OAS proteins. These amino acids were obtained by sequence and structural alignments against human and porcine OAS1. The residues are ordered so that analogous amino acids among the proteins are in the same position in the lists (e.g., Asp75, Asp408, and Asp816 are corresponding residues in OAS sequences).

#### Virtual screening

Virtual screenings against the donor and acceptor binding sites of OAS proteins were performed using the DrugDiscovery@TACC virtual portal [43], [44]. This web portal executed the Autodock Vina program on the “Lonestar” supercomputer at TACC (Texas Advanced Computing Center). The required input files were the target structures, the search spaces used for docking (Table S1), and the specification of the Virtual chemical library. While the protein structures in the “active-Mg^2+^-donor” state were used for screening on the acceptor binding site, the remaining targets were employed for screening on the donor binding site (Table 1). The chemical library used was a subset of the ZINC database termed the “ZINC (Lrg)” library [42], [43]. This library includes 642,769 small molecules with “clean, drug-like” constraints such as molecular weight between 150 and 500 Da, clogP ≤ 5, five or fewer hydrogen-bond donors, and ten or fewer hydrogen-bond acceptors. The analysis and visualization of the docking simulations were performed with UCSF Chimera 1.10.2, LIGPLOT+ 1.4.5 [45], and Discovery Studio Visualizer V17.2.0.16349 [46]

### Selection criteria

#### First selection filter

Two sets of selection criteria were established based on the binding site targeted on each protein state. Thus, ligands aiming at the donor binding site were filtered according to i) docking score lower than or equal to the positive control (Table S2); and ii) ability to interact with a minimum of six amino acids involved in the binding of the donor substrate, being at least two from the catalytic triad. Similarly, competitive inhibitors for the acceptor binding site were selected based on i) docking score lower than or equal to the positive control (Table S2); and ii) ability to interact with at least six amino acids implicated in the binding of the acceptor substrate. Any attractive contact within 3.7 Â was considered as an interaction if it was predicted by more than one analysis tool (UCSF Chimera 1.10.2, LIGPLOT+ 1.4.5, and Discovery Studio Visualizer v17.2.0.16349). Moreover, interactions with the catalytic triad included either direct contact with the aspartic acids or indirect interactions through Mg^2+^ ions.

#### Second selection filter

After selecting candidate molecules for each protein state, the ligands were docked at the remaining activation states within the same enzyme. Subsequently, the potential inhibitors were selected based on i) docking score lower than or equal to the positive control (Table S2); ii) ability to interact—in all protein states—with at least two amino acids from the catalytic triad, and two amino acids in charge of placing the donor substrate; and iii) ability to interact with a minimum of four amino acids implicated in positioning the acceptor substrate. Finally, to overcome the stochasticity behind docking calculations, all screening experiments were repeated five times; only candidate inhibitors found in all repetitions were considered for further analysis.

#### Toxicity and structure similarity

Using the OSIRIS Property Explorer [47], the candidate inhibitors were filtered based on the absence of risk alerts of mutagenicity or tumorigenicity. The final list of potential inhibitors was subjected to a structure similarity analysis utilizing the ChemMine Tools with a Tanimoto coefficient cutoff of 0.6 [48].

## Results

### Structural modeling and sequence alignment

To consider the activation mechanism of OAS proteins during the search for inhibitors of the 2-5A synthesis, homology models were constructed for the active and inactive states of human OAS1, OAS2, and OAS3. Five models preserving the characteristic folding of the active and inactive conformations were obtained with confidence scores (C-score) ranging from 1.5 to 1.79. The C-score, which usually goes from −5 to 2, being 2 the highest confidence [34]–[36], evidences the good quality of the predicted structures. Moreover, all models reproduced the conformational changes caused by the binding of RNA, such as the β-floor sliding and the formation of a new α-helix in the active site (Figure 1)[49]. Given the above, the models obtained were suitable representatives of the different activation states required for OAS activity and can be used for downstream analysis.

The amino acids that might be critical for the activity of OAS2 and OAS3 were determined based on human and porcine OAS1. By structural and sequence alignment, it was found that the catalytic triad and amino acids associated with the binding of the acceptor substrate were conserved among OAS1, OAS2, and OAS3 (Table 2 and Figure S2). Likewise, although slight differences were present at the donor site, all proteins preserved about 80% of the residues needed to position the donor substrate. The variations observed at OAS donor sites comprise the change of glutamine by lysine at position 566 in OAS2, and tyrosine by histidine at position 970 in OAS3 (Table 2 and Figure S2).

### Virtual screening and candidate inhibitor selection

Virtual screenings against the two ATP binding sites of OAS1, OAS2, and OAS3, considering four different activation states for each protein, led to the identification of molecules that could block OAS substrate sites at any protein state. A total of 12,000 compounds (1,000 molecules per activation state) were observed occupying, partially or entirely, the donor or acceptor binding site of OAS. No molecules were common to all the proteins; instead, most of the ligands were unique to specific states. After a filtering process (see Methodology: Selection criteria), it was found that a total of 454 compounds for OAS1, 672 compounds for OAS2, and 523 compounds for OAS3, could obstruct the binding of ATP in at least one of the activation states of the proteins. Notably, 15 molecules for OAS1,19 molecules for OAS2, and 7 molecules for OAS3 docked at all ATP binding sites, regardless of the activation state of the enzyme. These 41 compounds were subjected to a toxicity analysis as they represented the most likely competitive inhibitors of OAS proteins. Four candidates were discarded because of the presence of chemical fragments that could be associated with tumorigenesis or mutagenesis. The final 37 potential inhibitors of OAS enzymes are presented in Tables S4, S5, and S6.

### Protein-ligand interactions

Most of the activation states of OAS proteins take place at the donor binding site of the enzyme [12]. Consequently, the ability to interact with this region in the inactive, active, and active-Mg^2+^ states of the proteins was considered as an essential feature during the inhibitor selection. To identify characteristics associated with this property, the intermolecular contacts between the 37 candidate molecules and the OAS donor sites were analyzed using the interaction score method (Figure 2). As a result, it was observed that most of the compounds could interact with the same residues independently of the activation state of the enzyme. Specifically, OAS1 exhibited a consistent interaction profile where the amino acids Asp75, Asp77, Gln229, Gln194, and Tyr230 displayed the highest interactions score as they were in permanent contact with the inhibitors. Similarly, although OAS2 and OAS3 did not present a uniform pattern of interactions, the amino acids analogous to those described in OAS1 (except Gln194) also interacted with the ligands in the three activation states of these proteins (Figure 2).

**Figure 2.**
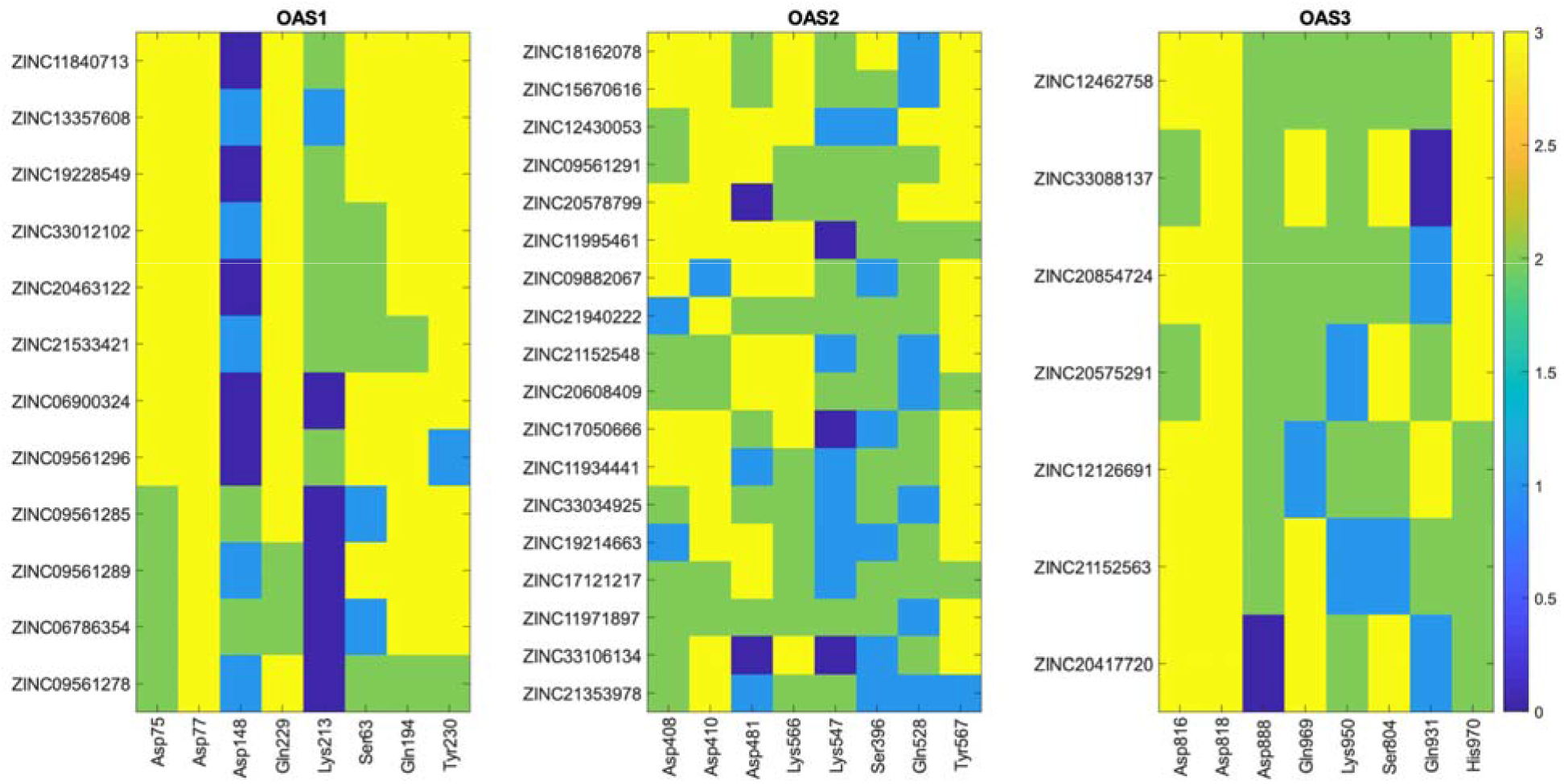
Overall contacts between potential OAS inhibitors and OAS donor sites. The colormap represents the number of OAS activation states where a candidate inhibitor interacted with an amino acid involved in the binding of the donor substrate. Only attractive contacts within 3.7À were considered as an interaction. The color code and scores are assigned as follows: score three (yellow) corresponds to interactions with the three activation states of the OAS donor site (inactive, active, and active-Mg^2+^); score two (green) is assigned to interactions presented in two activation states; score one (light blue) refers to interactions in one activation state; and score zero (dark blue) indicates no interaction. On the y-axes are presented the ZINC codes of the candidate molecules, sorted from the highest to the lowest total interaction score (see Tables S4, S5, and S6). On the x-axes are shown the residues involved in the binding of the donor substrate. These amino acids are ordered so that analogous residues among the three proteins are in the same position across the x-axes (e.g., Asp75, Asp408, and Asp816 are corresponding residues in OAS seguences).

To determine the predominant intermolecular contacts between the potential inhibitors and important amino acids for OAS activity, the frequency of five types of protein-ligand interactions was calculated. Thus, at OAS donor sites, although weak electrostatic contacts were dominant, strong non-covalent interactions were mainly established with the residues bearing the highest interaction scores (Figure 3A and Tables S4, S5, and S6). Accordingly, Asp75, Asp77, Gln229, and Tyr230 in OAS1 and their equivalents in OAS2 and OAS3 interacted commonly by anion-π interactions, hydrogen bonds, and π-π interactions. Remarkably, the non-conserved residues, such as Lys566 in OAS2 and His970 in OAS3, did not follow the previous tendency and showed a high frequency of alkyl-π contacts and hydrogen bonds, respectively. Regarding the acceptor binding site, similar interactions were observed between analogous residues, being electrostatic contacts and alkyl-π interactions the most frequent protein-ligand interactions (Figure 3.B).

**Figure 3.**
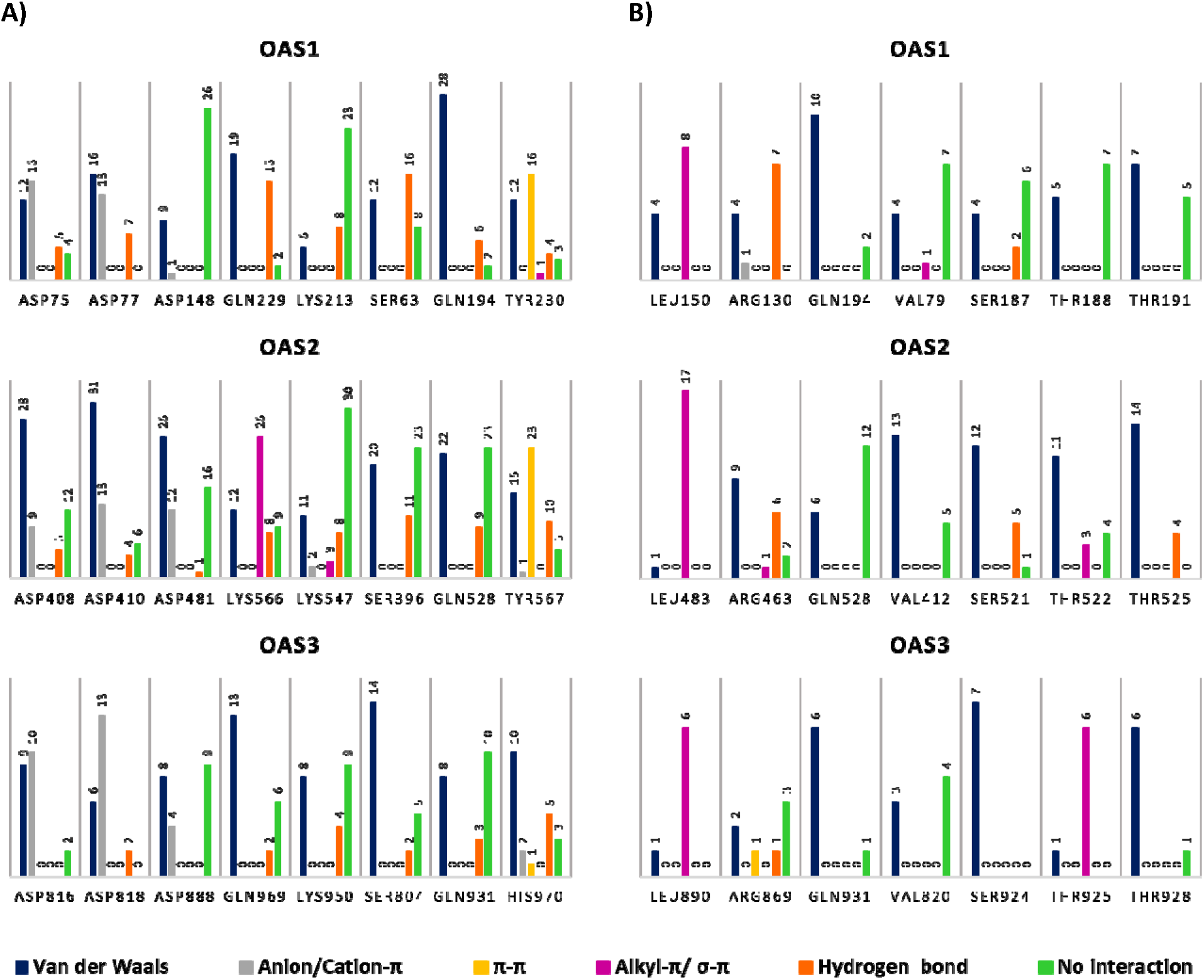
Frequency histogram of protein-ligand interactions between critical amino acids for OAS activity and potential OAS inhibitors. **A.** Protein-ligand interactions at OAS donor sites, considering three activation states for each protein (inactive, active, and active-Mg^2+^). **B.** Protein-ligand interactions at OAS acceptor sites. On the top of the bars is shown the frequency of each type of interaction, and on the x-axes are presented the residues involved in the binding of the donor and the acceptor substrates. In dark blue are shown Van der Waals contacts, in gray anion/cation-πeffects, in yellow π-π interactions, in magenta alkyl-π and σπ effects, in orange hydrogen bonds, and in green no interaction.

### Chemical structure similarity

Given the shared pattern of interactions across OAS1, OAS2, and OAS3, a structural similarity analysis was conducted among the potential inhibitors to identify scaffolds that could promote this behavior. As a consequence, two sets of highly similar molecules were identified for OAS1 (Figure 4), and small chemical fragments were found common to several compounds for OAS1, OAS2, and OAS3 (Table S3).

**Figure 4.**
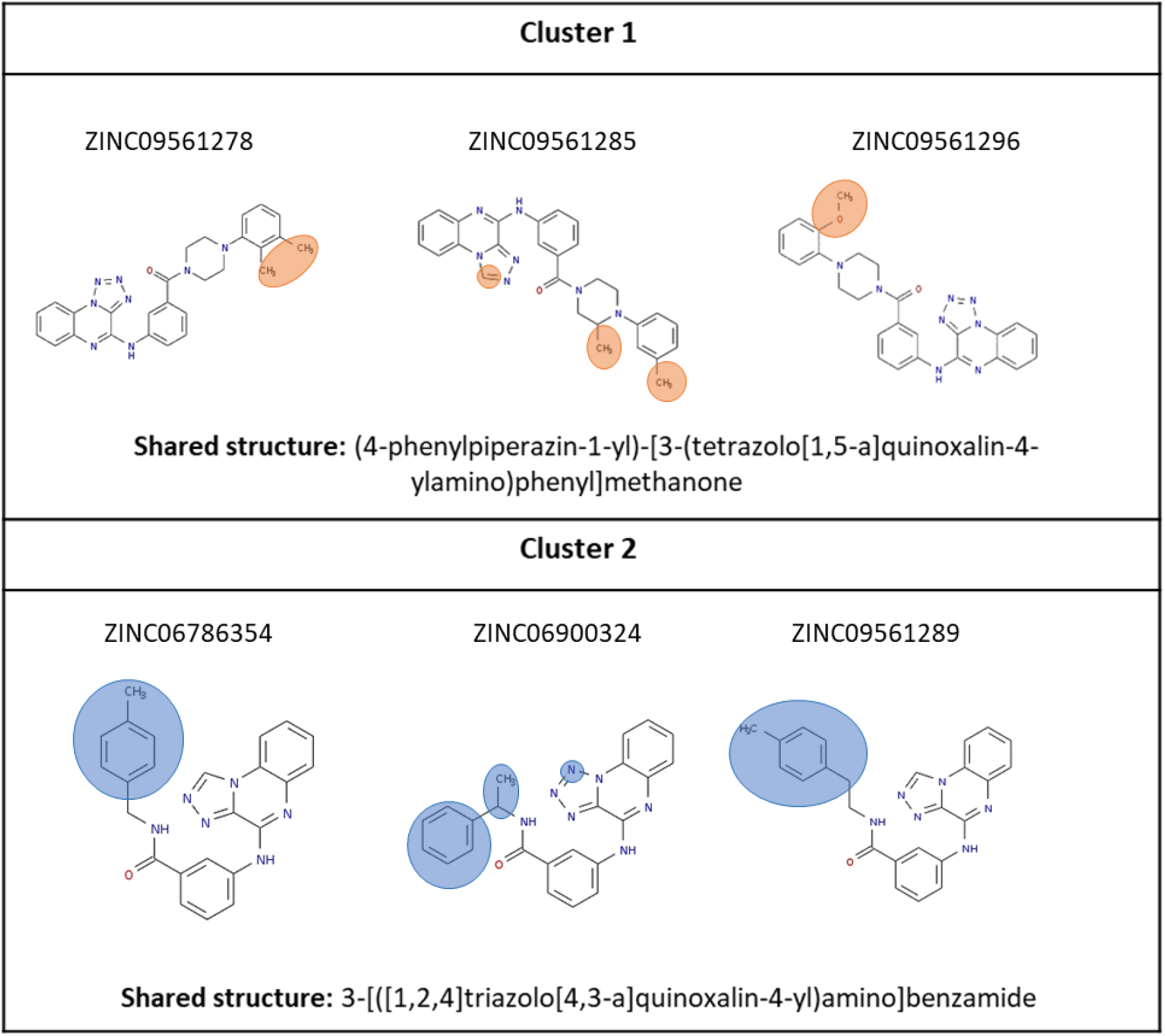
Shared structures among potential inhibitors for OAS1. The scaffolds were identified using ChemMine Tools with a Tanimoto coefficient cutoff of 0.6 [48]. In orange and blue ovals are highlighted the differences among the molecules.

The molecules containing shared structures were subjected to a detailed analysis of protein-ligand interactions to recognize intermolecular contacts associated with each chemical fragment. Although there was no consistency between interactions promoted by small chemical groups (Table S3), the scaffolds presented in Figure 4 did evidence a general interaction profile among the different activation states of OAS1. In this regarding, at the donor binding site of OAS1, the tetrazolo[1,5-a]quinoxaline ring (cluster 1), and the [1,2,4]triazolo[4,3-a]quinoxaline ring (cluster 2) established π-anion interactions with Asp75 and Asp77, and electrostatic contacts or hydrogen bonds with Gln229 (Figure 5, 6, and S4). Similarly, at the acceptor binding site, the [1,2,4]triazolo[4,3-a]quinoxaline ring contacted Leu150 through π-alkyl interactions while the tetrazolo[1,5-a]quinoxaline ring tended to form hydrogen bonds with Arg130 (Figure S5). On the other hand, even though the phenylpiperazinyl group in cluster 1 exerted different types of contacts with OAS1, this chemical fragment was usually positioned towards amino acids involved in the binding of the acceptor substrate (Figure 5 and 6). This latter characteristic was also observed in other compounds containing a phenylpiperazinyl group, such as ZINC33034925 (OAS2), and ZINC20417720 (OAS3) (Figure 3S).

**Figure 5.**
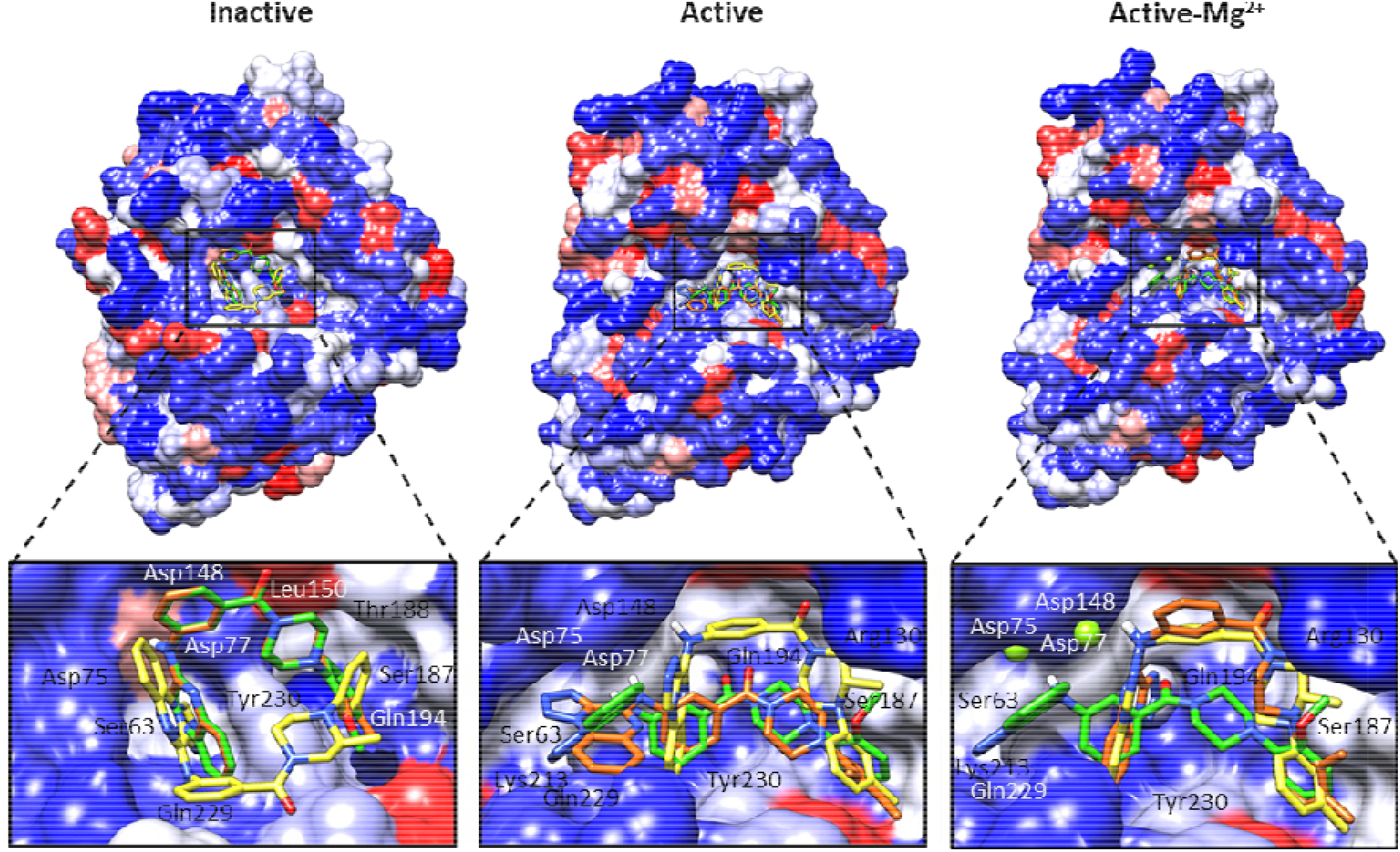
Donor binding site of OAS1 in complex with potential inhibitors sharing the scaffold (4-phenylpiperazin-1-yl)-[3-(tetrazolo[1,5-a]guinoxalin-4-ylamino)phenyl]methanone. The molecules presented belong to cluster 1 in Figure 4. In yellow is displayed ZINC09561285, in orange is presented ZINC09561278, in green is shown ZINC09561296, and green spheres represent Mg^2+^ atoms.

**Figure 6.**
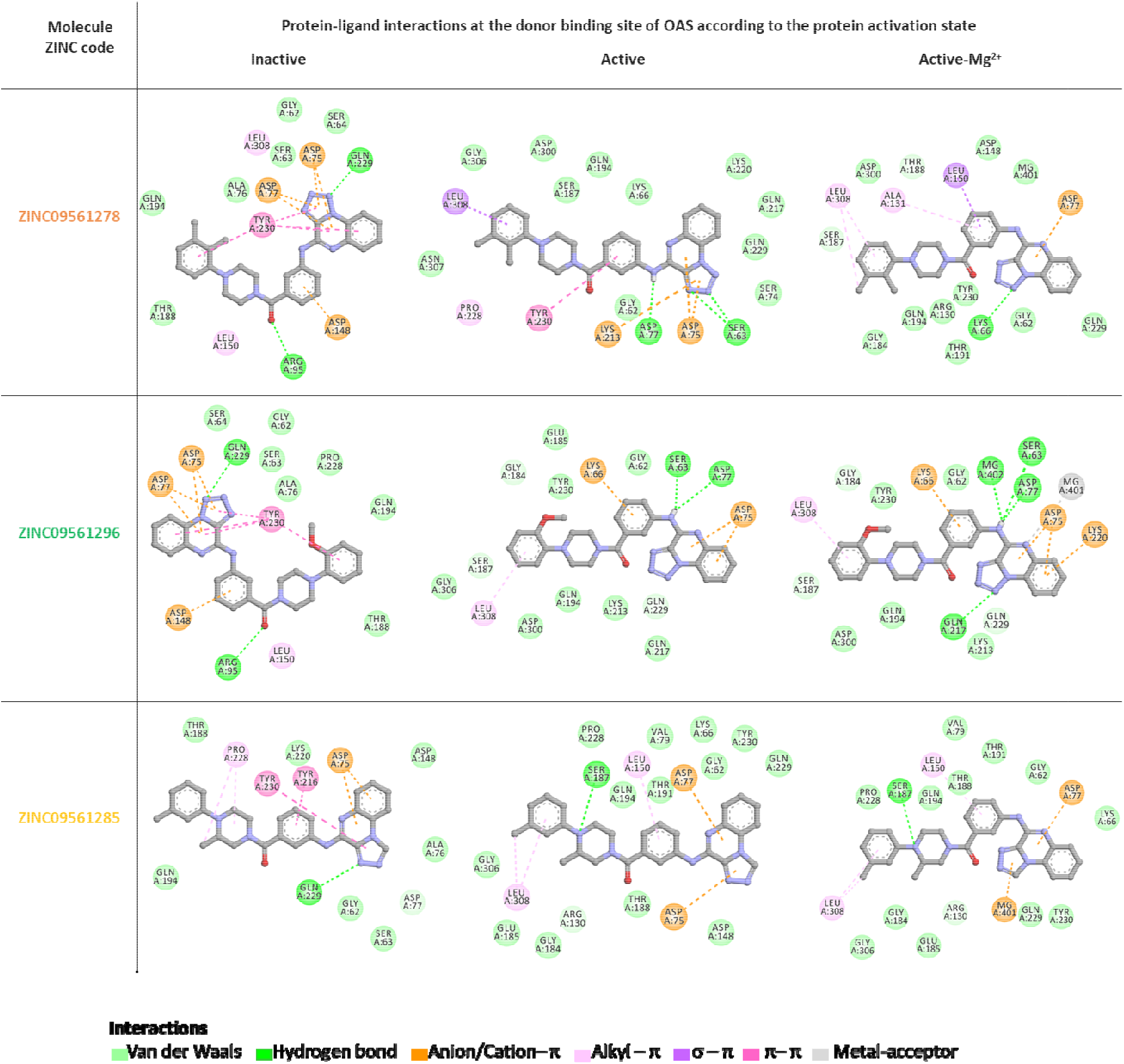
Predicted intermolecular interactions between the donor binding site of OAS1 and potential inhibitors sharing the scaffold (4-phenylpiperazin-1-yl)-[3-(tetrazolo[l,5-a]guinoxalin-4-ylamino)phenyl]methanone. The molecules presented belong to cluster 1 in Figure 4. The ZINC codes of the compounds are colored according to their binding modes in Figure 5. The carbon atoms in the molecules are shown in gray, nitrogen atoms are presented in blue, and oxygen atoms are indicated in red. The color code used to represent the intermolecular interactions is: In light green are presented Van der Waals interactions, in bright green hydrogen bonds, in orange anion-π or cation-π effects, in pink or purple hydrophobic interactions, in magenta π-π effects, and in gray metal-acceptor contacts.

## Discussion

The OAS family of proteins have been involved in a variety of immunoregulatory functions, promoting the progression of diseases such as cancer [22], autoimmune and autoinflammatory disorders [17]–[21], and several infectious diseases [23], [24]. These detrimental outcomes are generally the result of a continuous positive feedback loop between the activation of OAS and the production of type I interferon [28]. The possibility of disrupting this cycle of excessive inflammatory responses by the inhibition of OAS proteins places these enzymes as a new target for immune modulation.

In nature, many viruses counteract OAS activity by sequestering their dsRNA or through modification or degradation of the 2-5A messenger [50]–[53]. Moreover, zinc, copper, and iron ions have also demonstrated to impair OAS function by altering the protein oligomerization [14]. Besides this, there is no evidence of other mechanisms or compounds that can interrupt OAS action. In this study, we propose an ATP-competitive inhibition as a potential strategy to impede the 2-5A synthesis. As a result, we have identified 37 small molecules that can dock at the two ATP binding sites of OAS1, OAS2, or OAS3, with lower energy scores than the natural complexes. Furthermore, the compounds here presented can adjust to the dynamic behavior of these proteins and interact with their substrate sites independently of the activation state of the enzyme.

Despite the inherent variations among OAS proteins, a general pattern of protein-ligand contacts was observed at the donor binding site of OAS1, OAS2, and OAS3. Particularly, the residues Asp75, Asp77, Gln229, and Tyr230 in OAS1, and their equivalents in OAS2 and OAS3 were in constant interaction with the candidate inhibitors (Figure 2). Notably, although this region undergoes several conformational changes, the stated amino acids seem to be stable as they keep their position throughout OAS activity. Even if our homology models might not capture the precise spatial location of these residues, likely, interactions with the “stable” regions of the binding site promote the docking of ligands at any activation state of the protein. Indeed, numerous strong non-covalent interactions (hydrogen bonds and π-effects) were observed between the steady amino acids and the potential inhibitors (Figure 3A).

Although distinct chemical groups contributed to a general pattern of protein-ligand interactions, certain molecular properties were common within the candidate inhibitors. Overall, it was observed that π-systems containing hydrogen bond acceptors (mostly heterocycles) could interact with OAS active site regardless of the activation state of the protein. In this respect, the π-clouds played a fundamental role in the binding of the inhibitors as they promoted π-anion interactions with the catalytic triad, π-cation contacts in the presence of Mg^2+^, and π-alky or π-π effects with residues neighboring the active site (Figure 6, and Figures S3-S5).

In summary, this work proposes 37 small drug-like molecules as potential ATP-competitive inhibitors of OAS proteins. Altogether, our results suggest that the candidate compounds can interrupt the formation of the 2-5A messengers by binding at the substrate sites of the active and inactive conformations of OAS proteins. Considering the influence of the OAS family in the development of inflammatory disorders, the ligands here identified represent potential alternatives for host-centric interventions aiming at modulating excessive inflammatory responses. As a future direction, experimental validation will be needed to determine the precise inhibitory potential of these compounds.

## Conclusion

The inhibition of the OAS family of proteins represents a host-centric strategy that could reduce the severity of a broad spectrum of diseases. In this study, we have identified 37 small drug-like compounds (12 molecules for OAS1, 18 molecules for OAS2, and 7 molecules for OAS3) that could exert a competitive inhibition at the donor and acceptor binding sites of OAS proteins. Several commonalities among the candidate inhibitors, such as general patterns of protein-ligand interactions as well as shared chemical fragments, are valuable findings that could be further explored in the rational design and optimization of general or selective OAS inhibitors. Overall, the molecules here presented offer a new set of candidates for in vitro testing and characterization as potential OAS inhibitors.

## Acknowledgments

This work was supported by the Defense Advanced Research Projects Agency and the US Army Research Office through the program Technologies for Host Resilience - Host Acute Models of Malaria to study Experimental Resilience (THoR’s HAMMER), DARPA contract #W911NF-16-C-0008, 2016-2019. Additionally, this project was also supported by the Voluntary Tuition Incentive Program for Research Grants funded by the Graduate School at the University of Georgia. The authors acknowledge the Texas Advanced Computing Center (TACC) at The University of Texas at Austin for providing High-performance computing (HPC) and the drug databases that have contributed to the research results reported within this paper. URL: http://www.tacc.utexas.edu

## Supplementary Information

**Figure SI.**
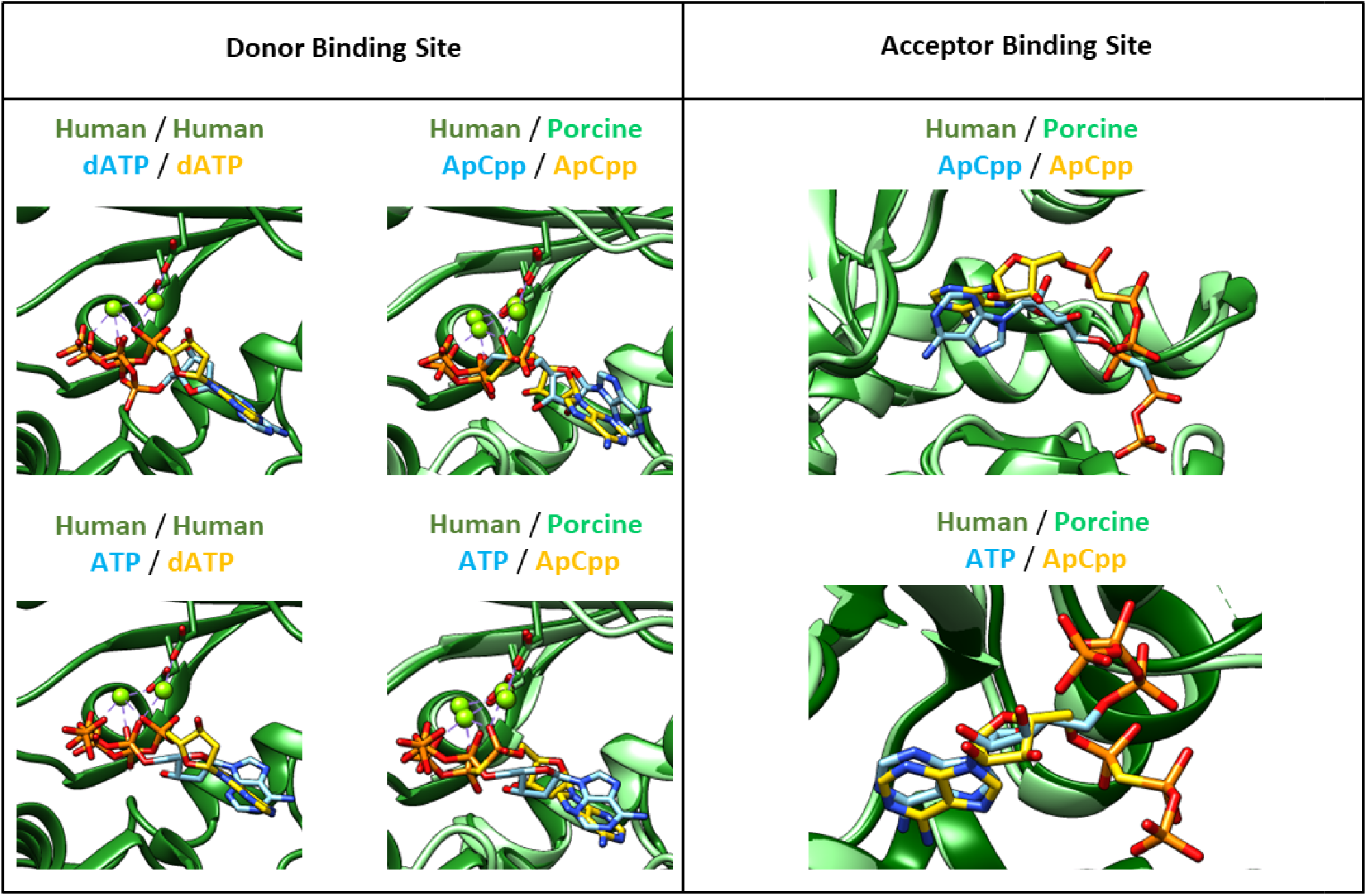
Docking results approximating the natural binding of OAS substrates. The proteins displayed corresponds to human OAS1 <u>4IG8</u> (dark green) and pOASl <u>4RWN</u> (light green). On the left are compared the binding modes of donor substrates on the OAS1 “Active-Mg^2+^” state. The ligands docked (dATP, ApCpp, and ATP) were contrasted against the corresponding human and porcine crystal structures. In yellow is presented the binding of the ligand in the crystallization and in blue are shown the docking results. Green spheres were used to display the Mg^2+^ ions. Since ATP has not been crystallized in complex with OAS, the binding mode of this molecule was compared against the binding of dATP and ApCpp. On the right are analyzed the binding modes of the acceptor substrates on the OAS1 “Active -Mg^2+^-donor” state. The compounds studied as acceptors were ApCpp and ATP.

**Figure S2.**
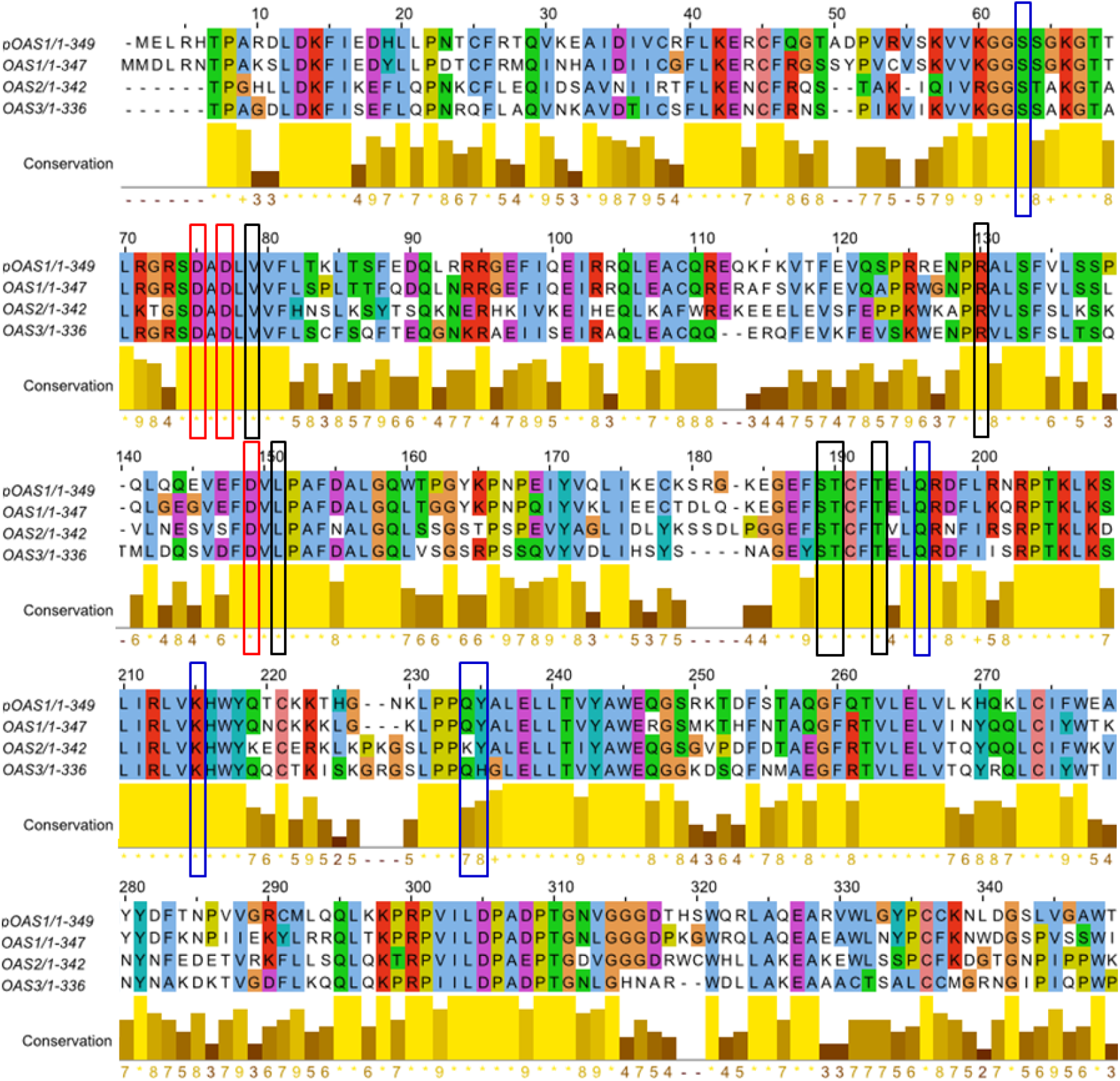
Sequence alignment between pOAS1 and human OAS1, OAS2, and OAS3. The color code used corresponds to the default Clustal X color scheme, which is assigned depending on the chemical properties of the amino acids. In red squares are highlighted the amino acids belonging to the catalytic triad, in blue squares are underlined the residues involved in positioning the donor substrate, and in black squares are emphasized the amino acids associated with the binding of the acceptor substrate. The numbering does not refer to a position in a specific sequence, but the position in the alignment.

**Figure S3.**
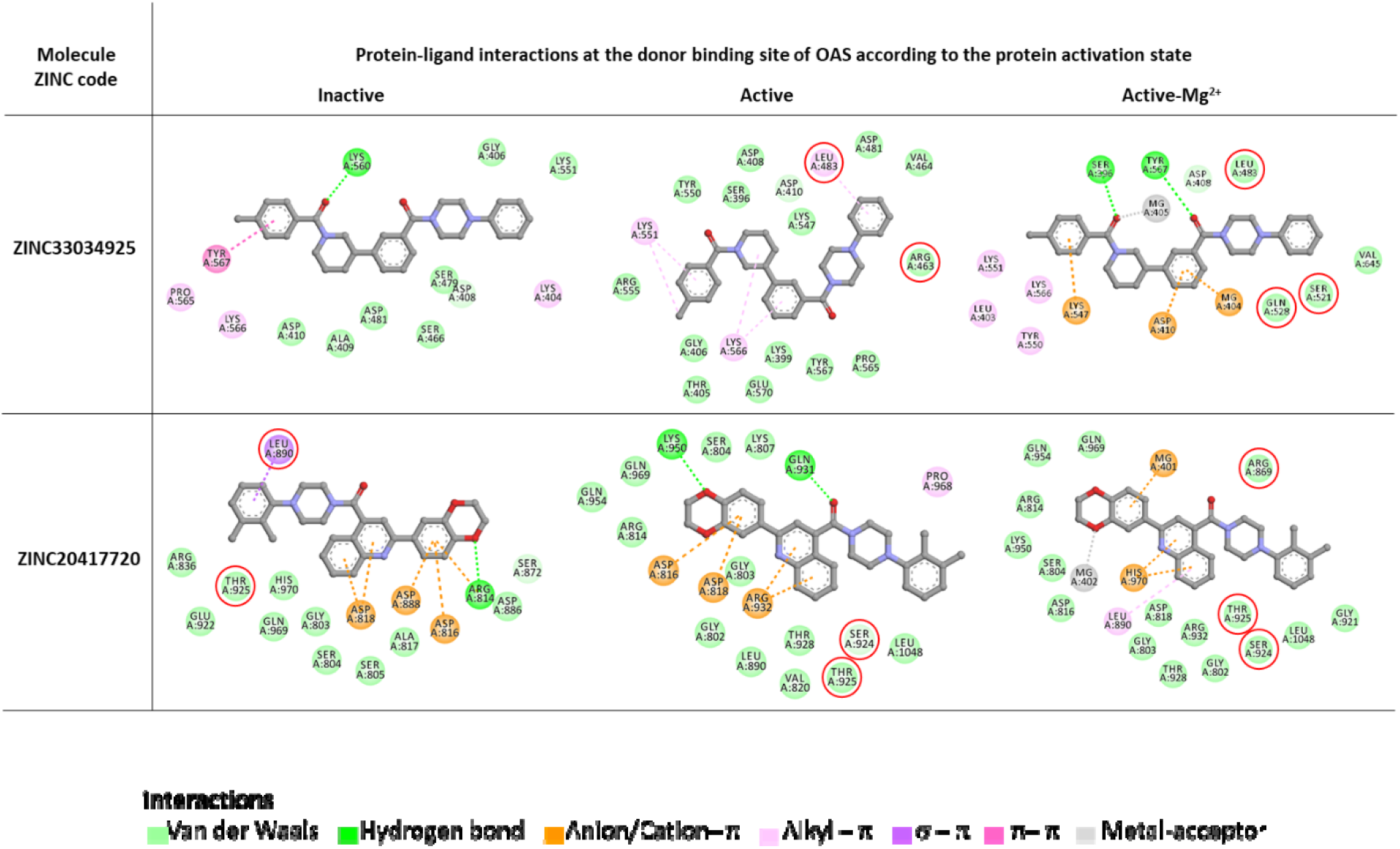
Protein-ligand interactions among OAS donor sites and candidate inhibitors containing a phenylpiperazinyl fragment. ZINC33034925 is docked on OAS2, and ZINC20417720 is interacting with OAS3. The phenylpiperazinyl group is usually observed in contact with residues involved in the binding of OAS acceptor substrates (red ovals). The carbon atoms of the ligands are shown in gray, nitrogen atoms are presented in blue, and oxygen atoms are indicated in red. The color code used to represent the intermolecular interactions is: In light green are presented Van der Waals interactions, in bright green hydrogen bonds, in orange anion-π or cation-π effects, in pink or purple hydrophobic interactions, in magenta π-π effects, and in gray metal-acceptor contacts.

**Figure S4.**
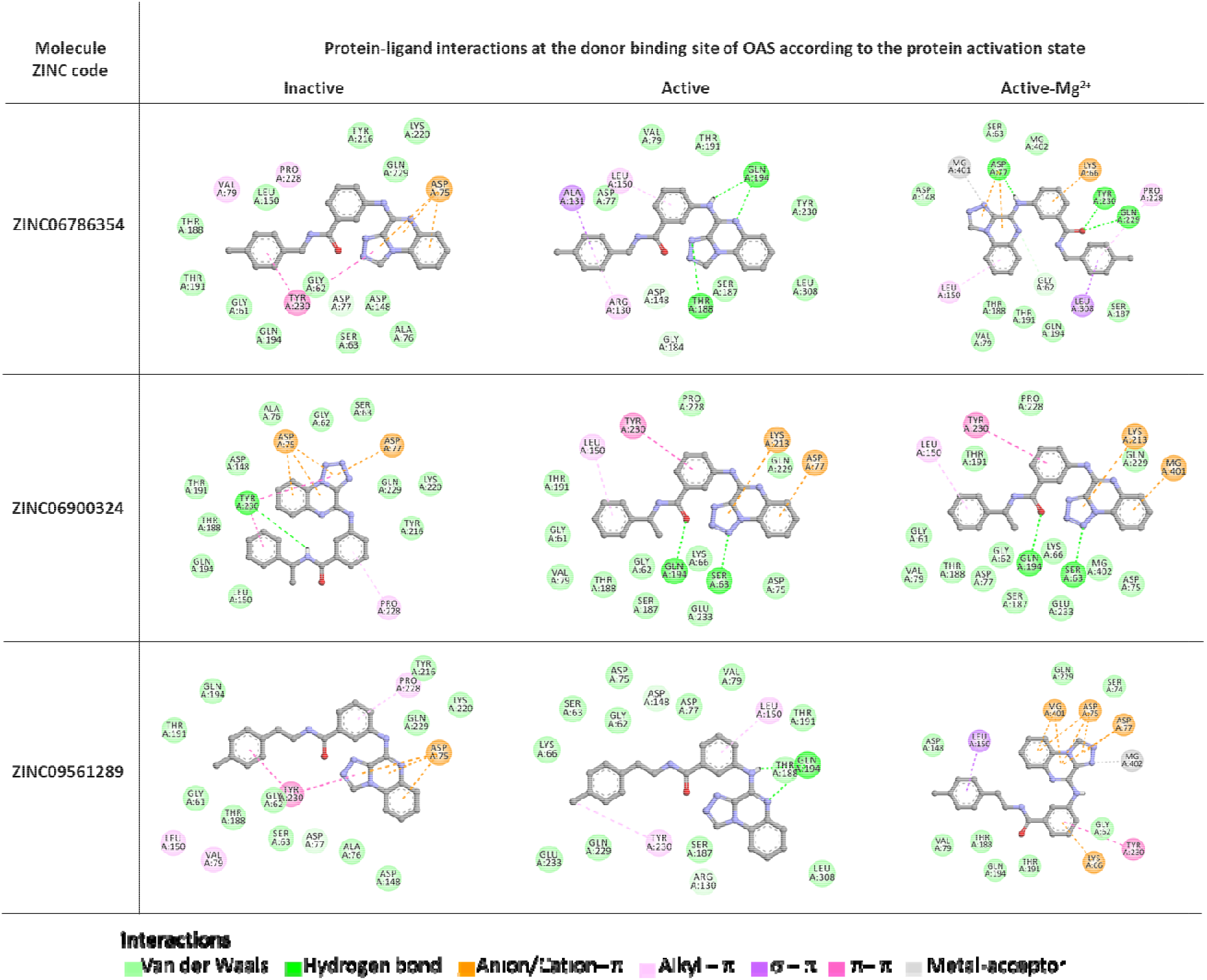
Intermolecular interactions between the donor binding site of OAS1 and candidate molecules sharing the scaffold 3-[([1,2,4]triazolo[4,3-a]guinoxalin-4-yl)amino]benzamide. The molecules presented belong to cluster 2 in Figure 4. The carbon atoms of the potential inhibitors are shown in gray, nitrogen atoms are presented in blue, and oxygen atoms are indicated in red. The color code used to represent the intermolecular interactions is: In light green are presented Van der Waals interactions, in bright green hydrogen bonds, in orange anion-π or cation-π effects, in pink or purple hydrophobic interactions, in magenta π-π effects, and in gray metal-acceptor contacts.

**Figure S5.**
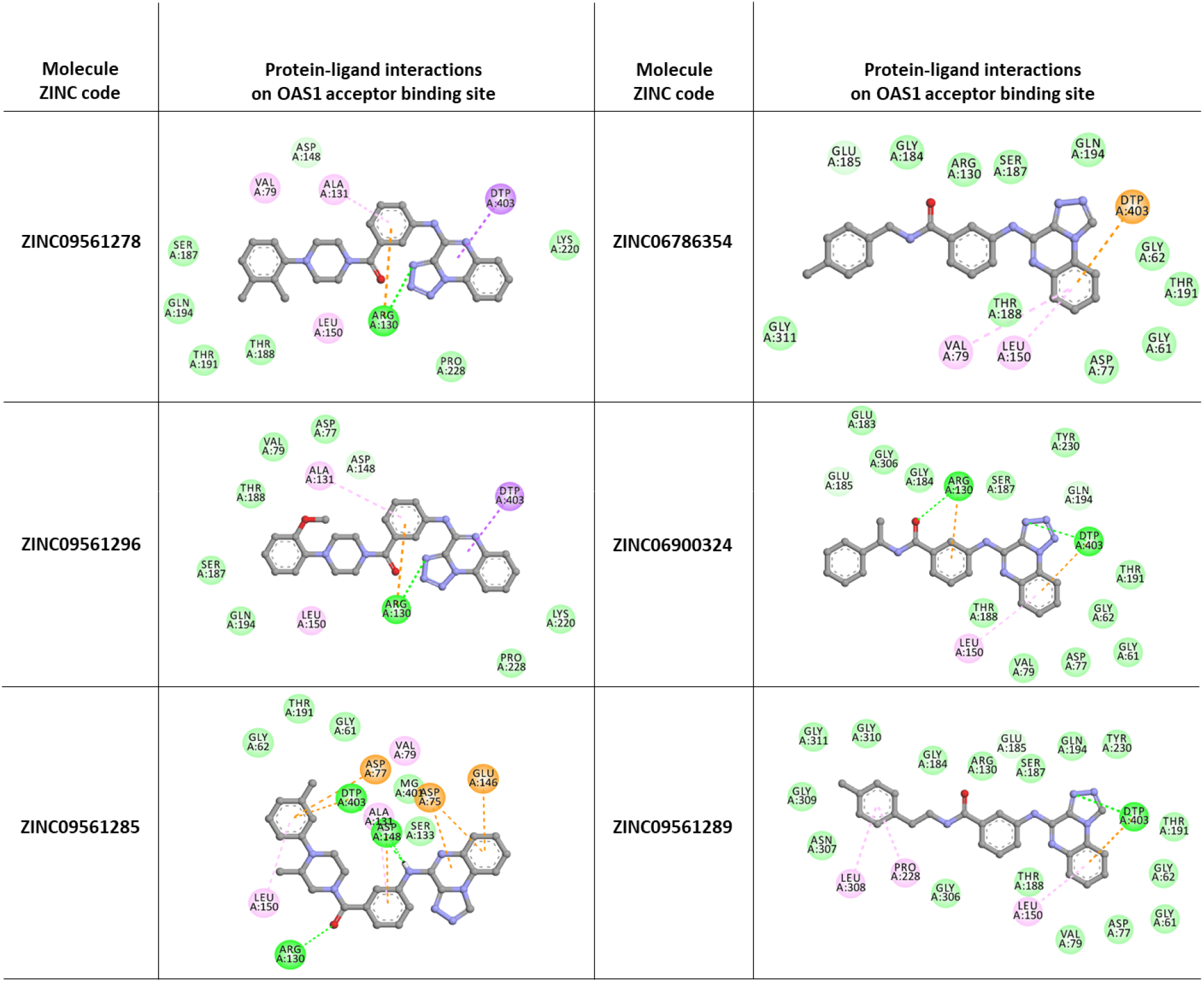
Intermolecular interactions between the acceptor binding site of OAS1 and candidate molecules with high structural similarity. Referring to Figure 4, on the left are presented molecules belonging to cluster 1, and on the right are shown molecules grouped on cluster 2. The carbon atoms of the potential inhibitors are shown in gray, nitrogen atoms are presented in blue, and oxygen atoms are indicated in red. The color code used to represent the intermolecular interactions is: In light green are presented Van der Waals interactions, in bright green hydrogen bonds, in orange anion-*π* or cation-π effects, in pink or purple hydrophobic interactions, in magenta π-π effects, and in gray metal-acceptor contacts.

**Table S1.**
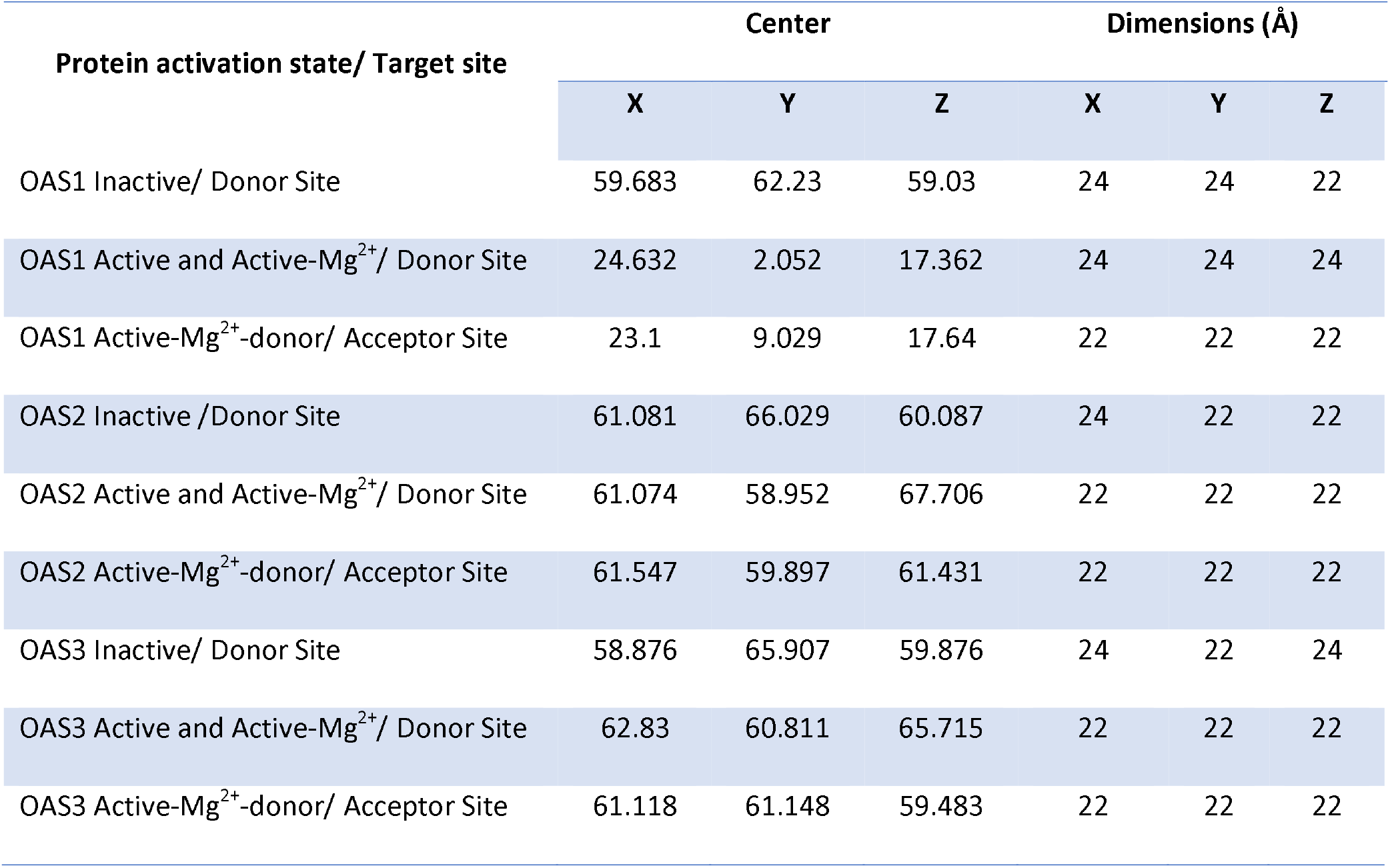
Search spaces used for virtual screening against donor and acceptor binding sites of human OAS proteins.

**Table S2.**
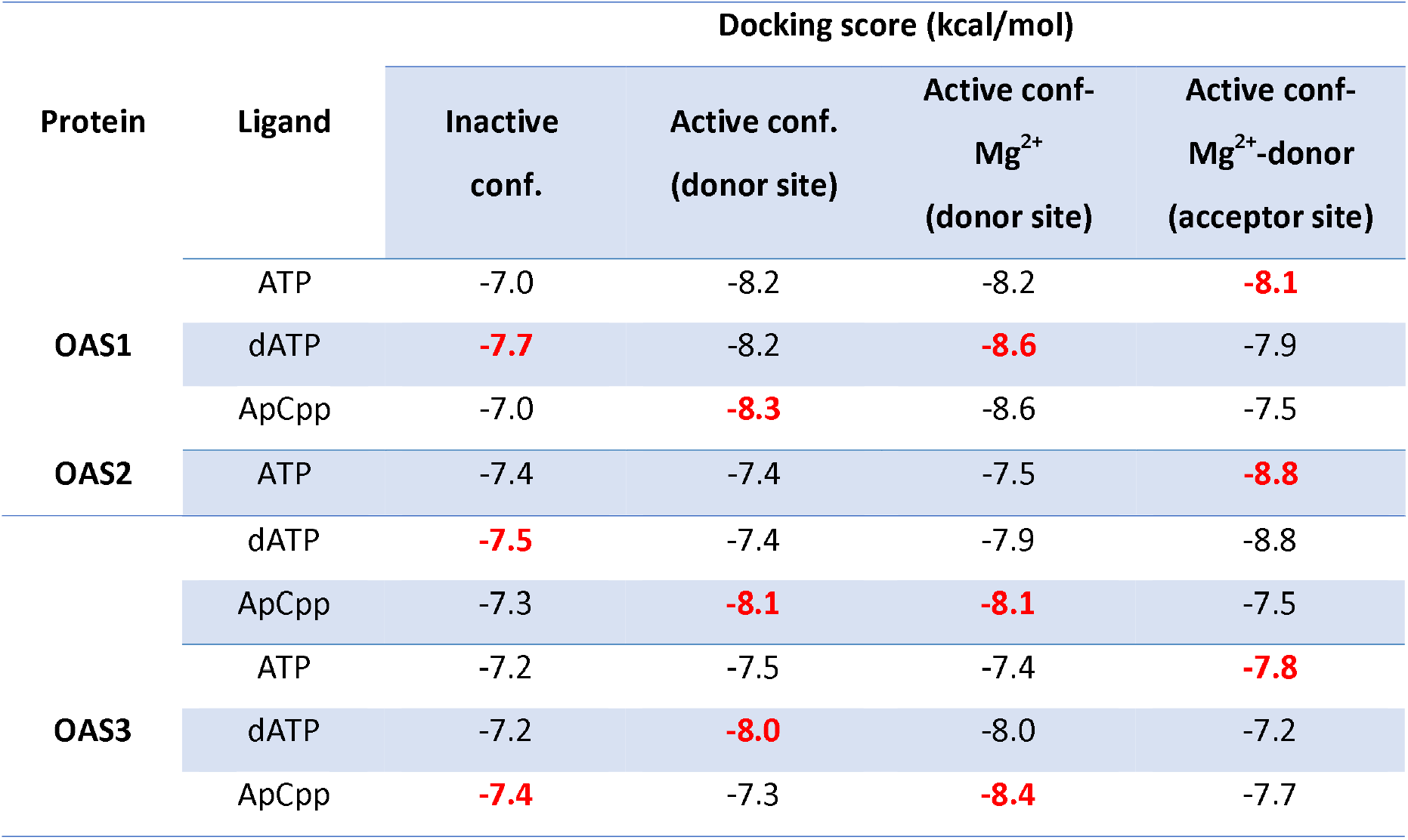
Docking scores of ATP and ATP analogs in complex with human OAS1, OAS2, and OAS3. In red are highlighted the docking scores used as positive controls during the candidate selection. These values correspond to the lowest docking score among the docking results with ATP, dATP, and ApCpp.

**Table S3.**
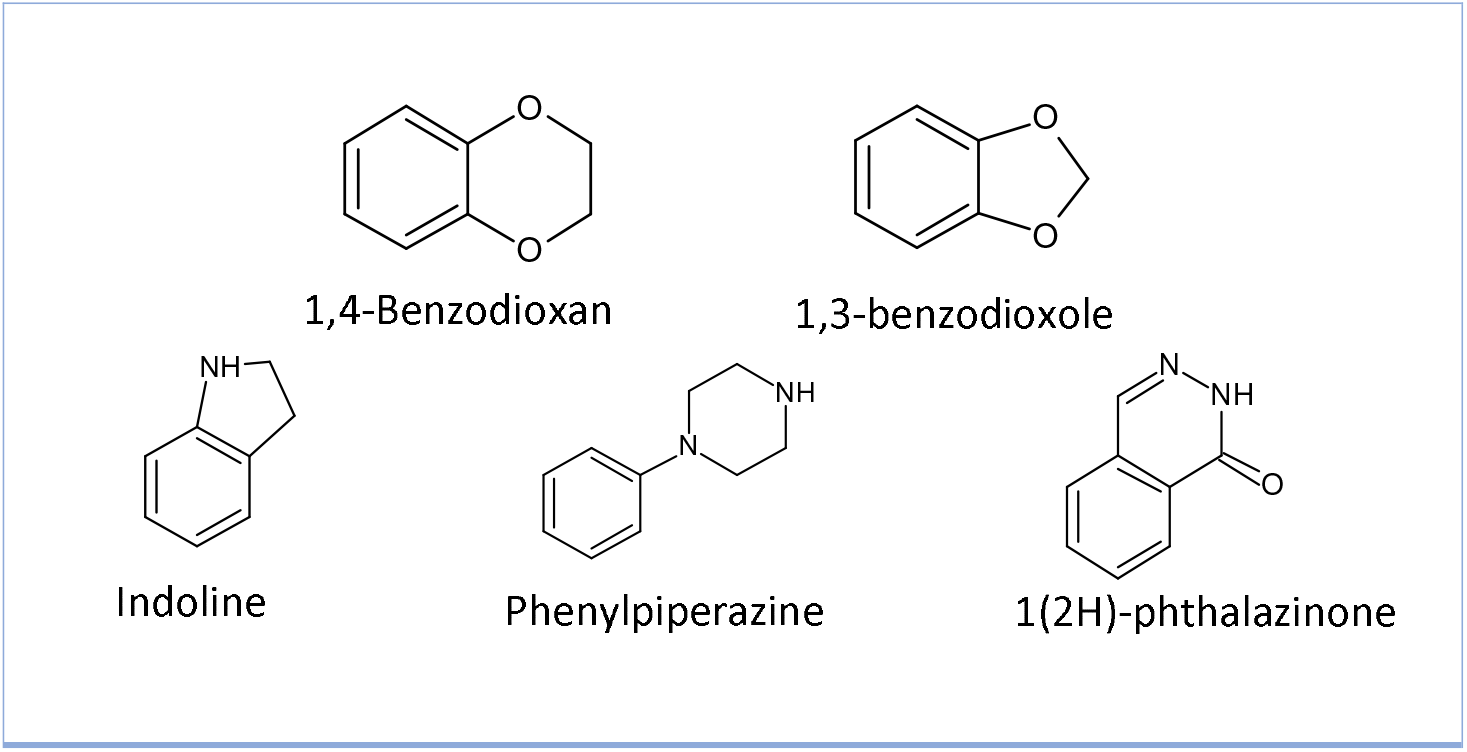
Common chemical groups among OAS potential inhibitors.

**Table S4.**
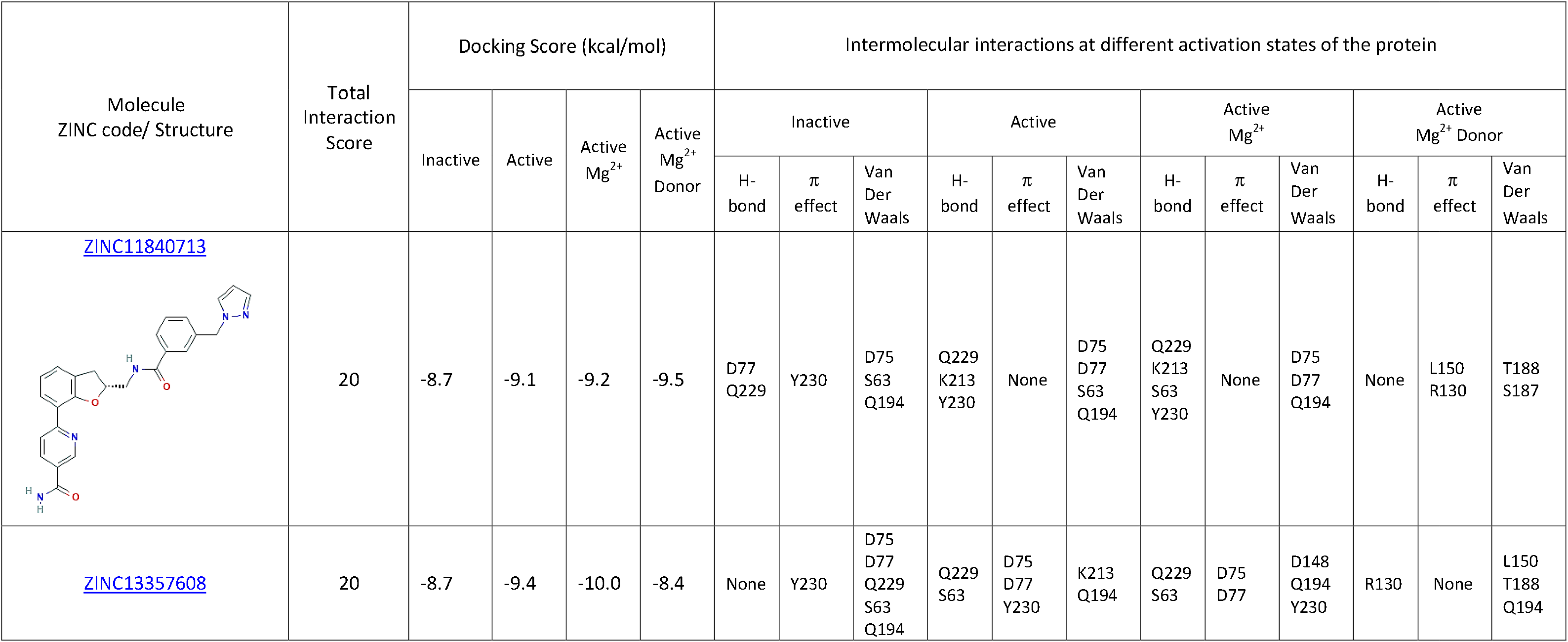
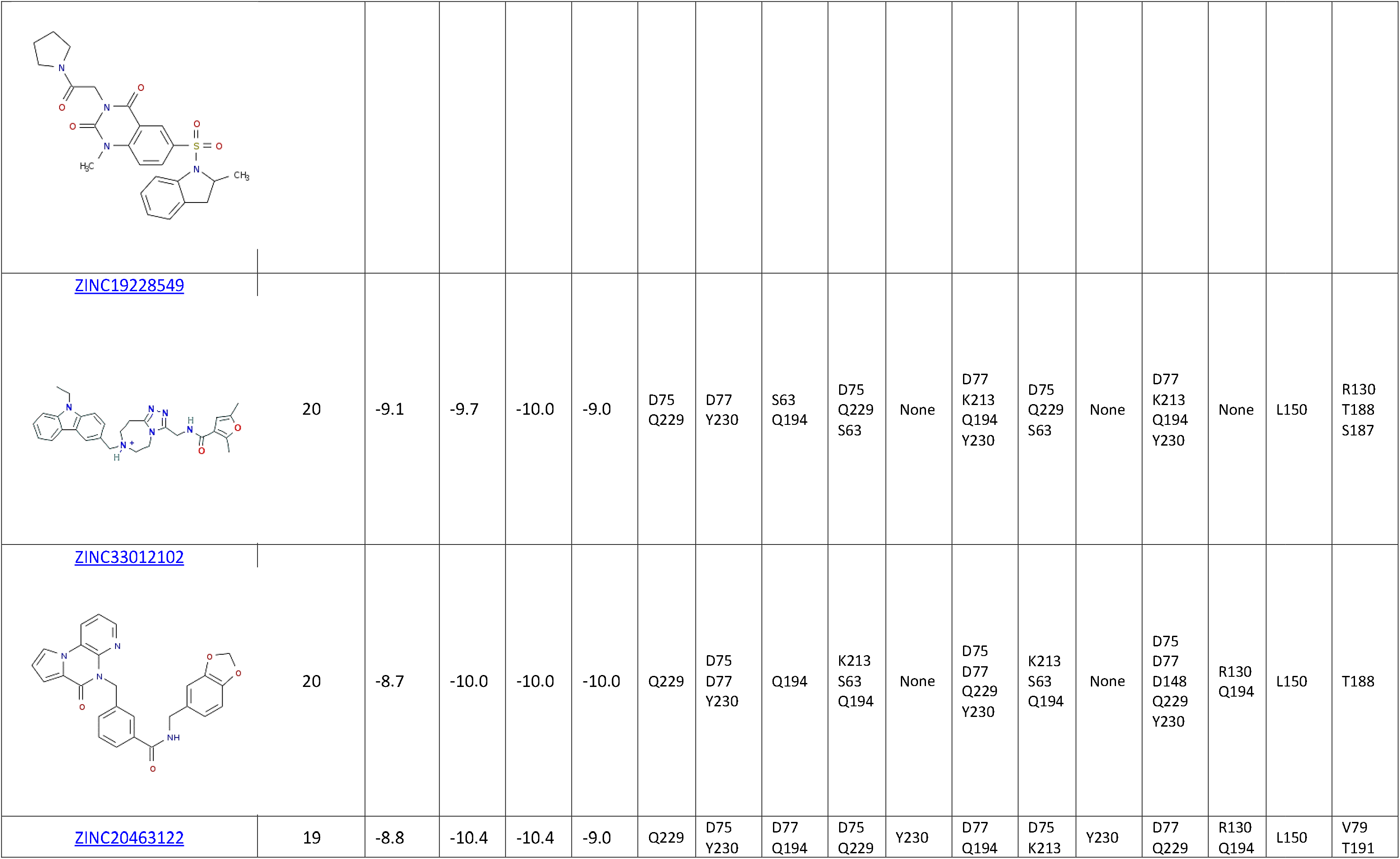
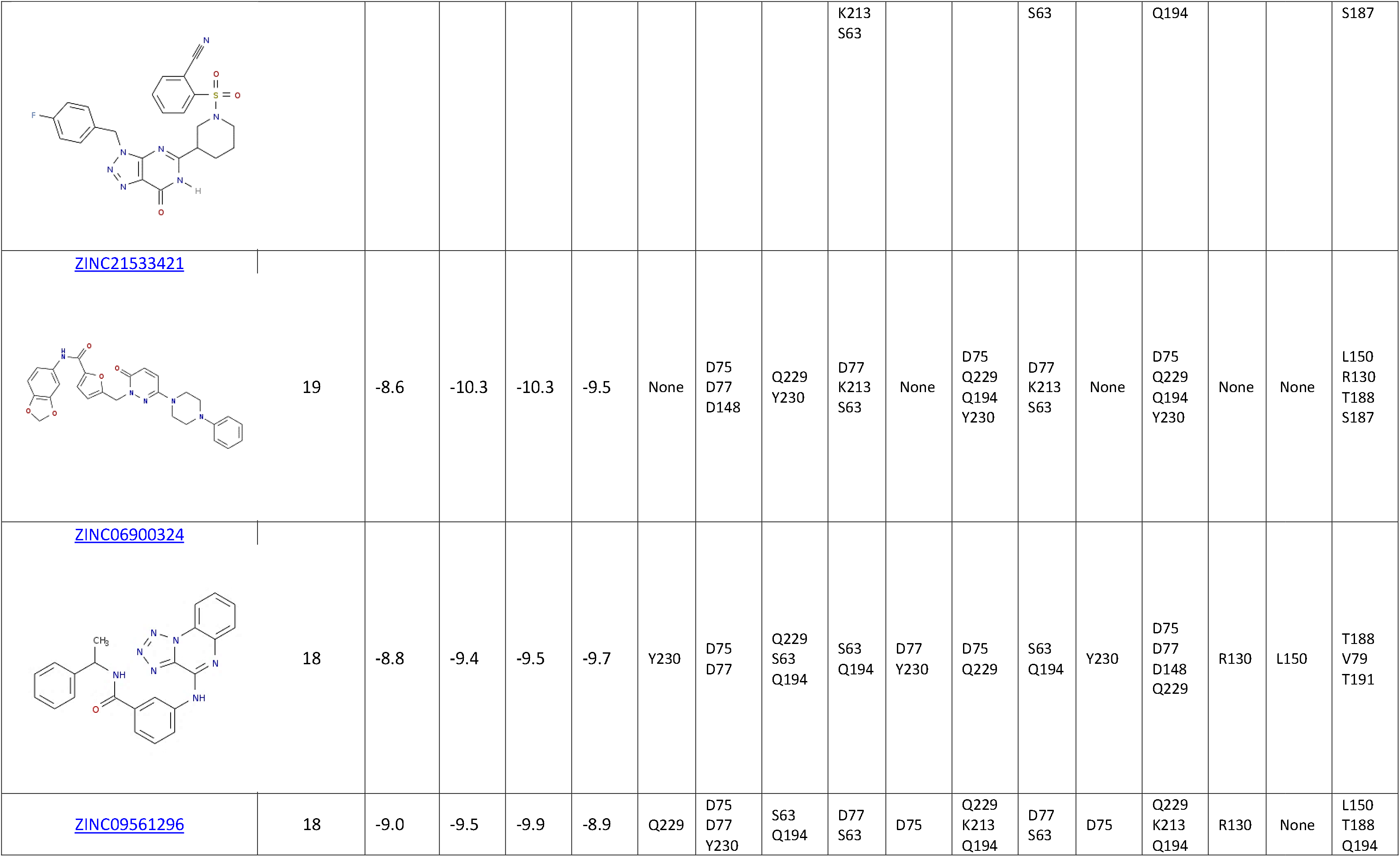
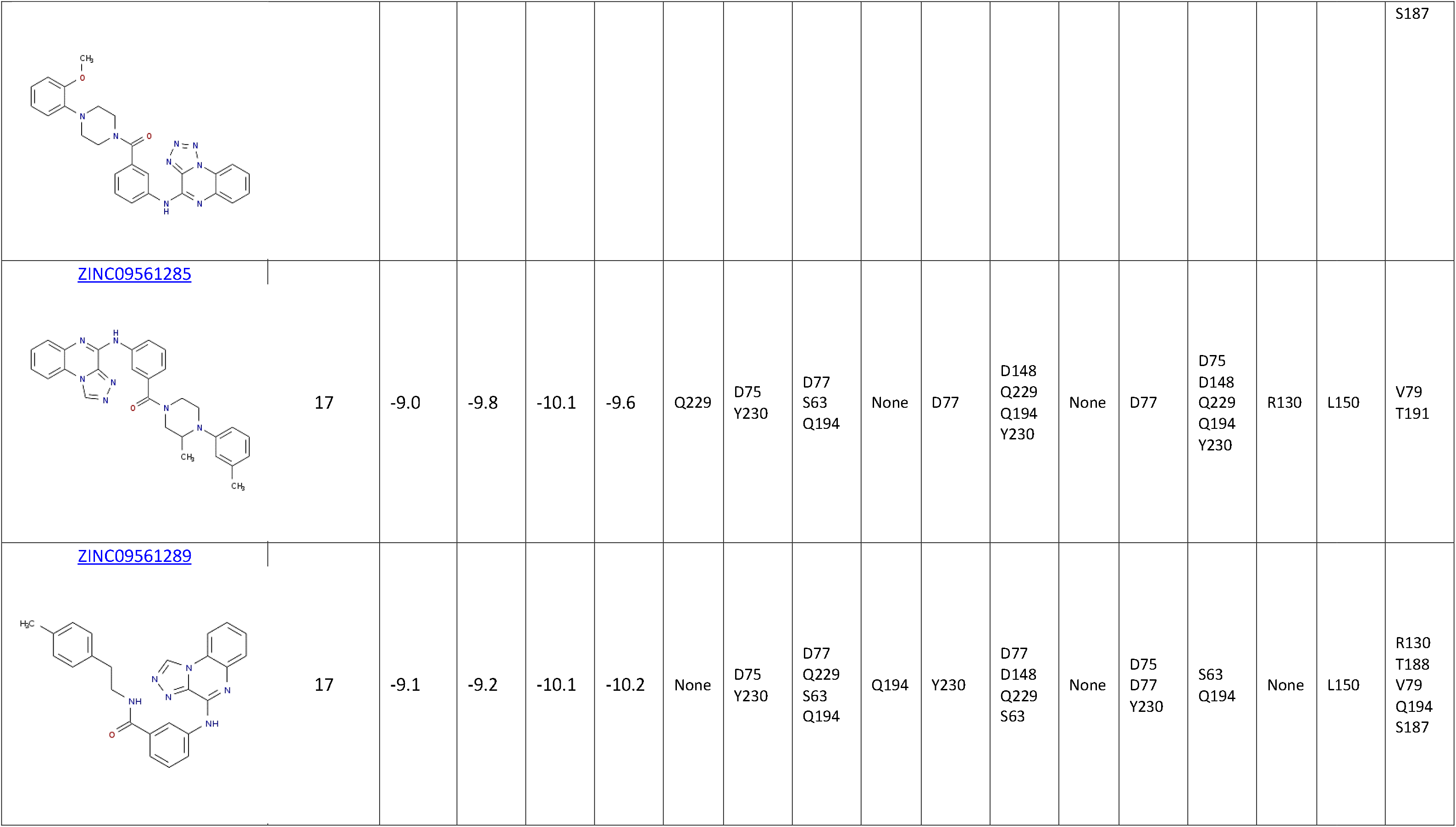
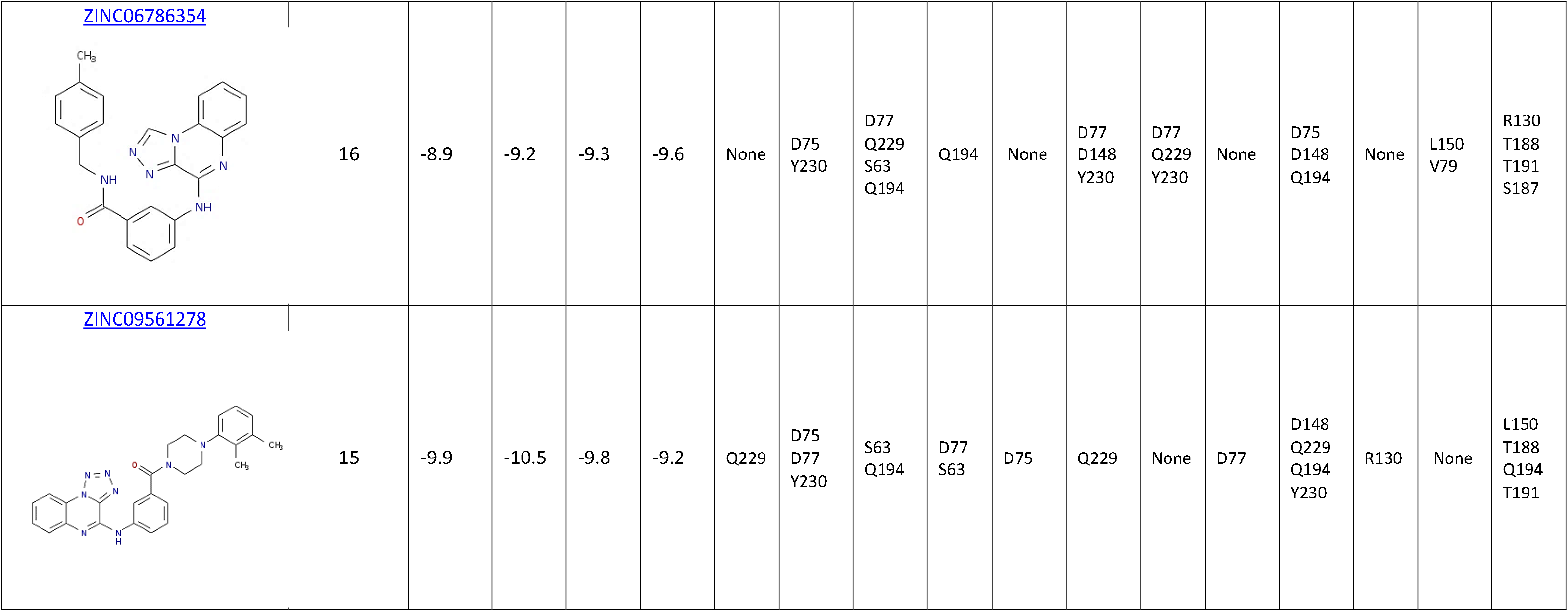
Information associated with potential inhibitors for OAS1. The total interaction score was calculated as the sum of the interaction scores per residue (Figure 2). The intermolecular interactions presented only consider amino acids involved in the binding of the donor or the acceptor substrate. Web pages with details about the candidate molecules and biological tests are attached to each ZINC code. This information was retrieved from [54].

**Table S5.**
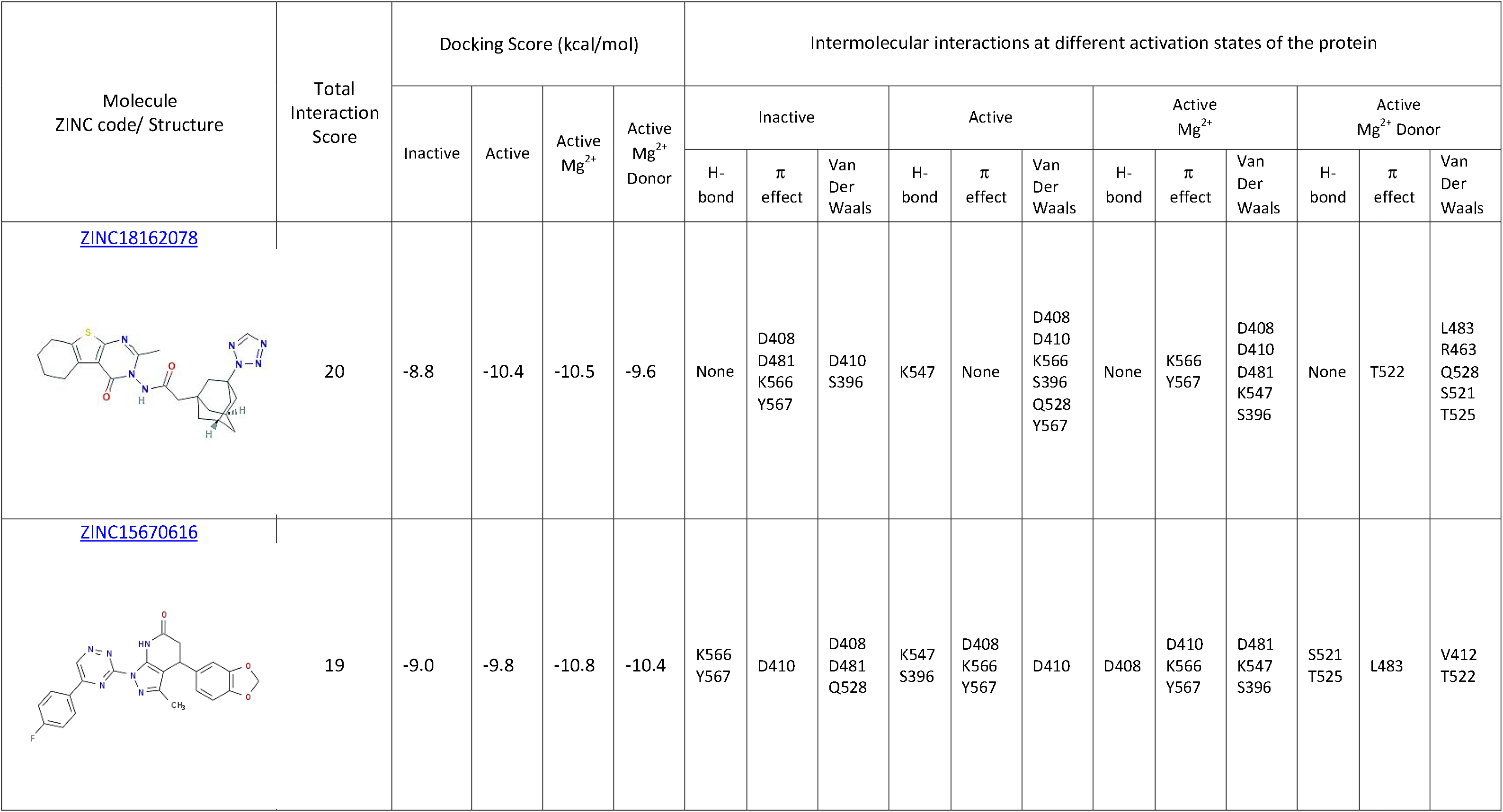
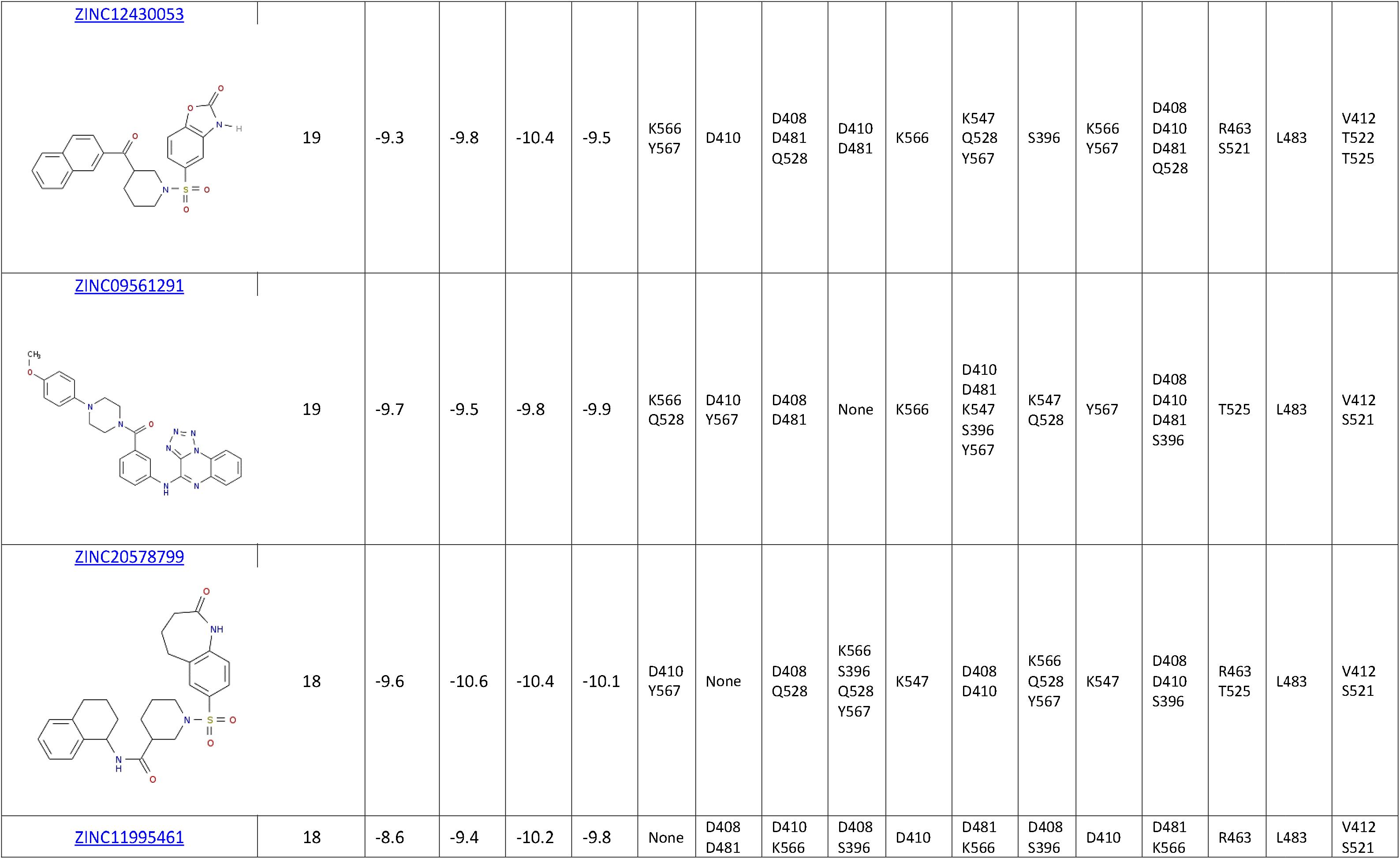
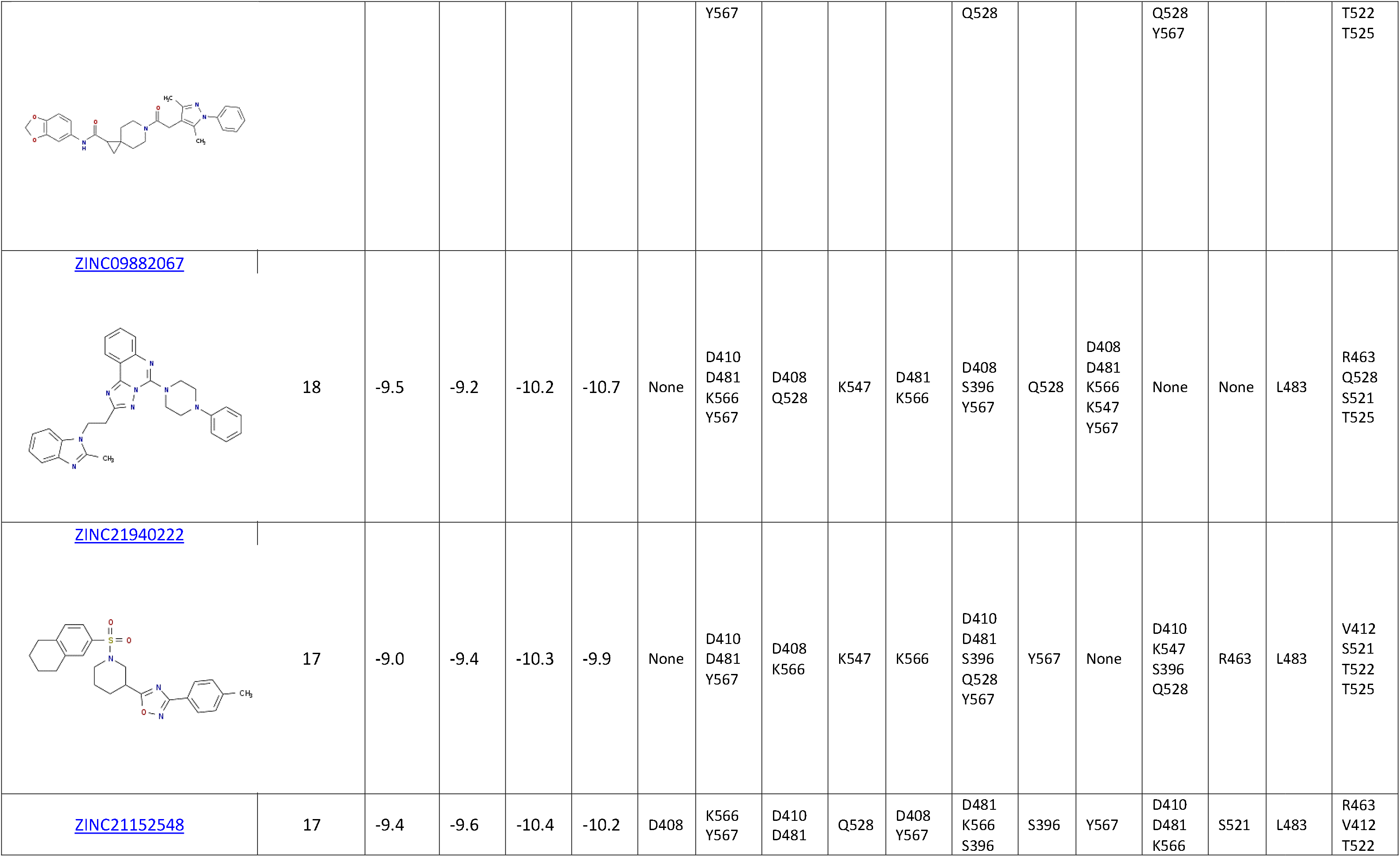
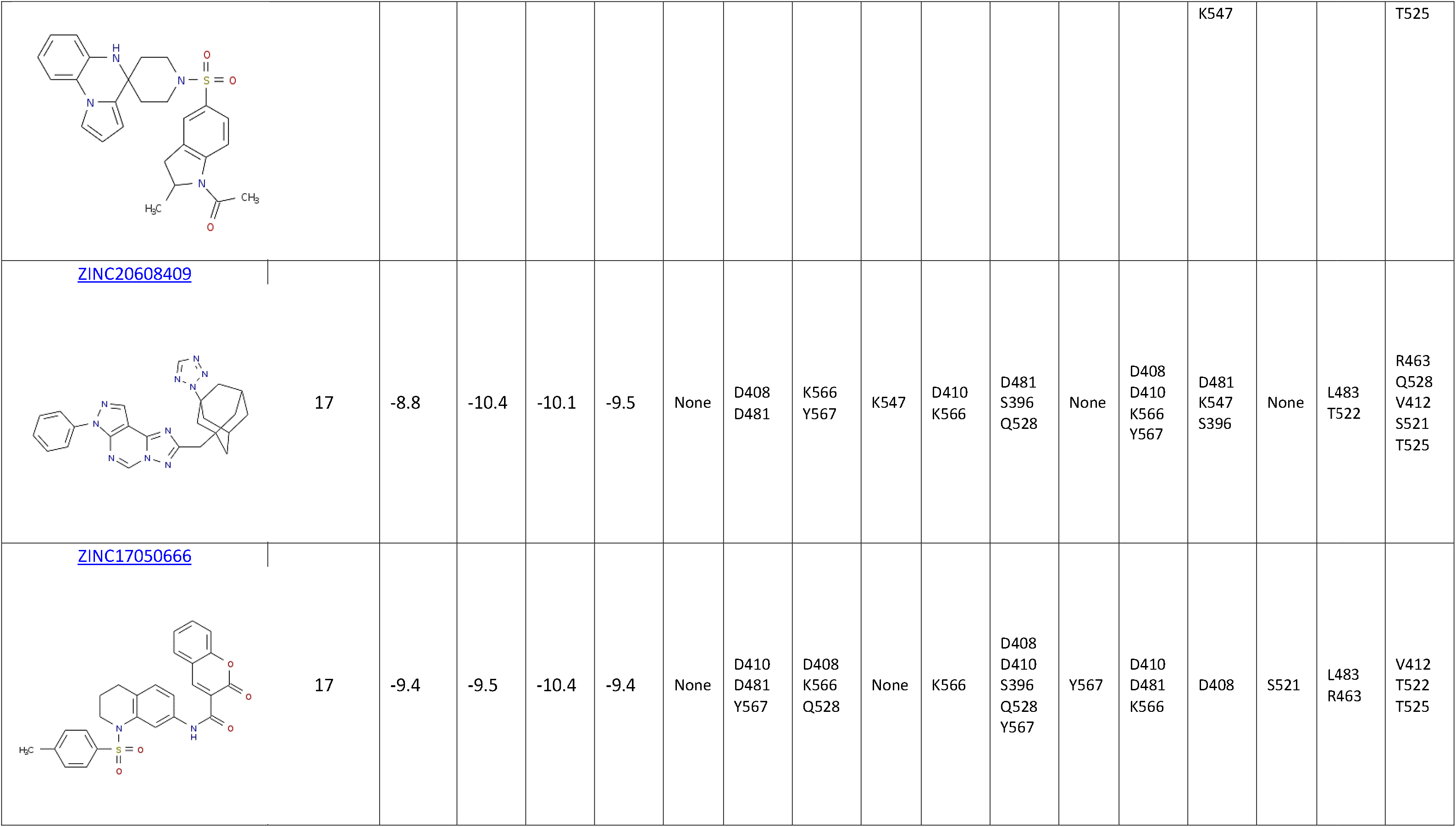
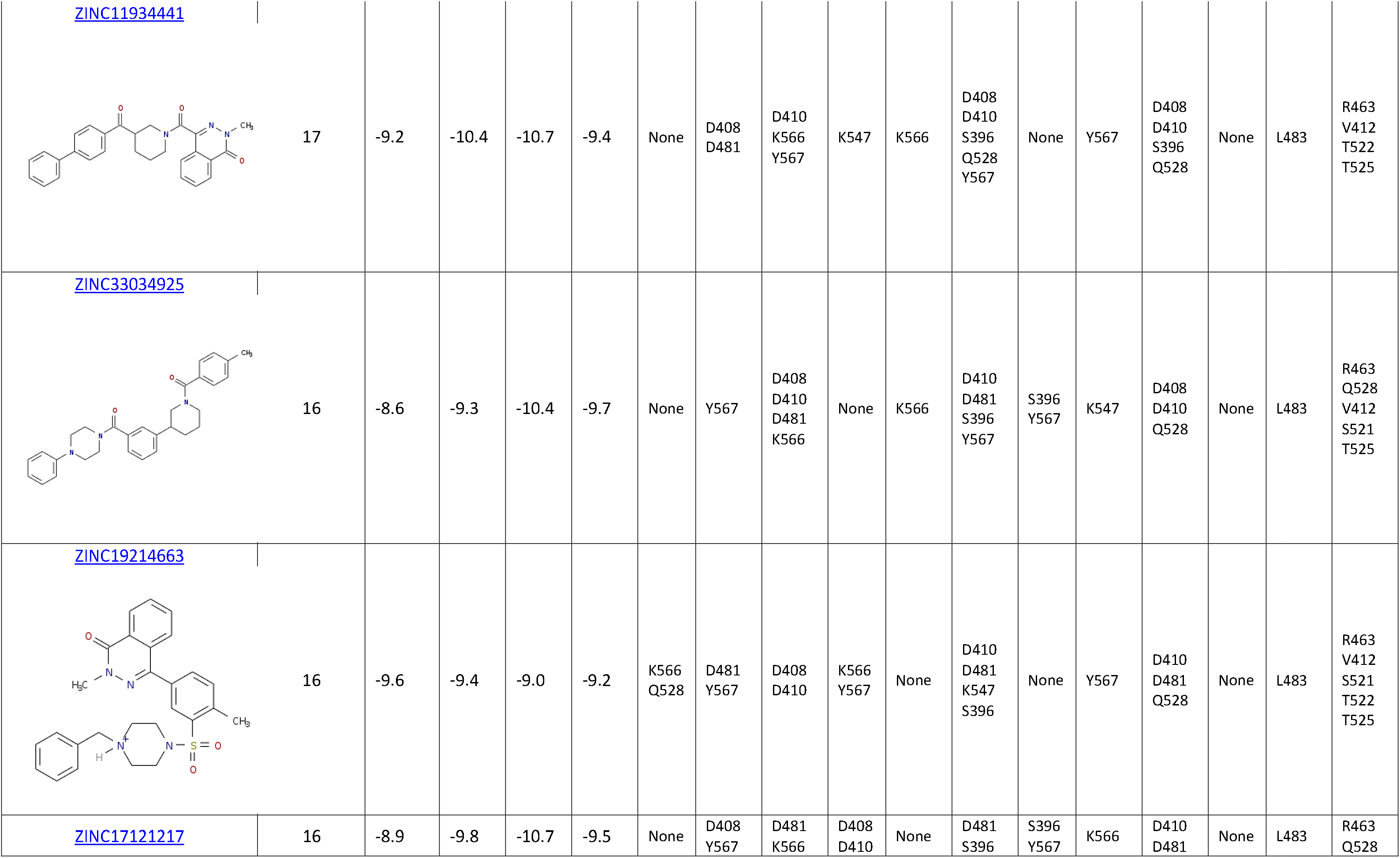
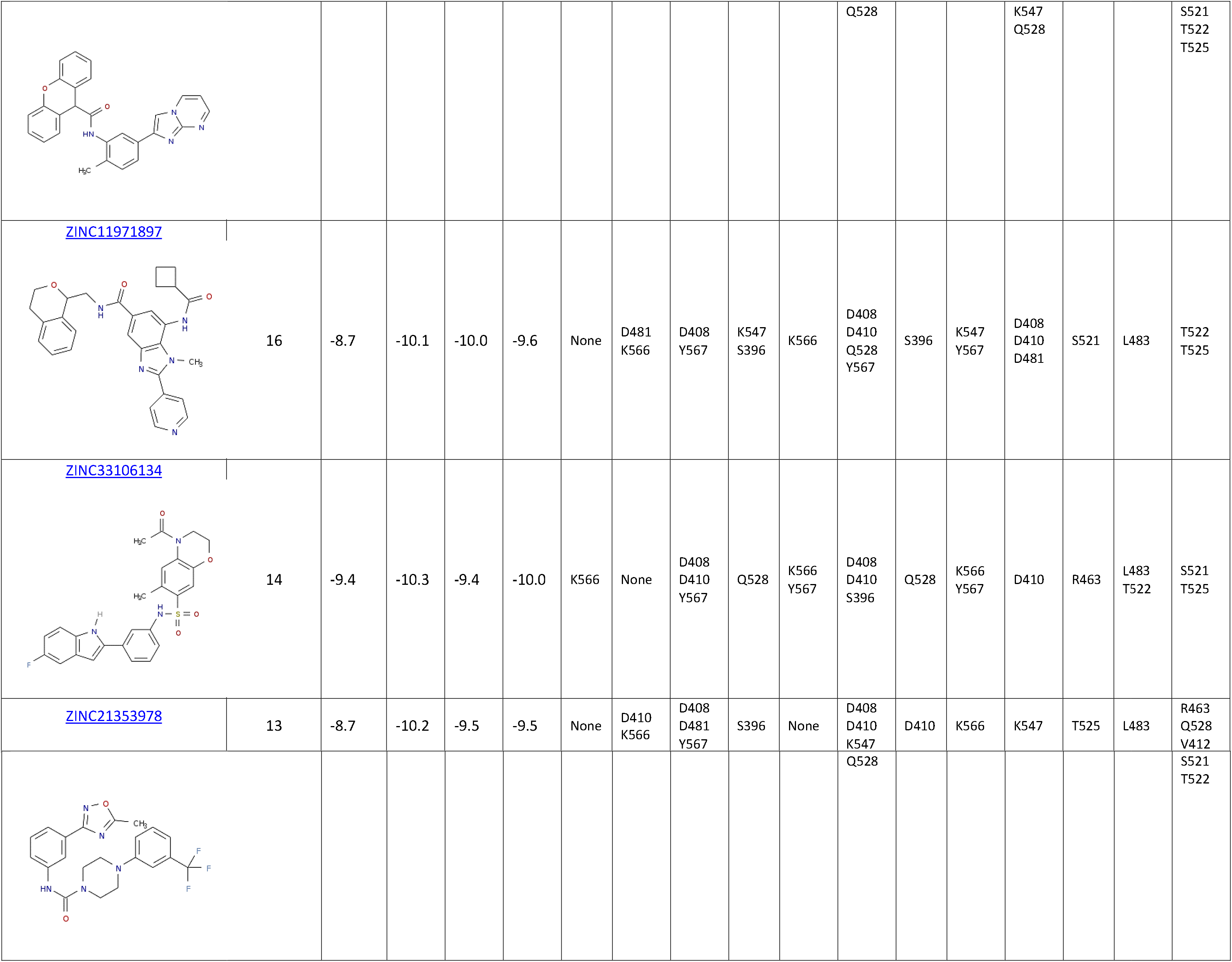
Information associated with potential inhibitors for OAS2. The total interaction score was calculated as the sum of the interaction scores per residue (Figure 2). The intermolecular interactions presented only consider amino acids involved in the binding of the donor or the acceptor substrate. Web pages with details about the candidate molecules and biological tests are attached to each ZINC code. This information was retrieved from [54].

**Table S6.**
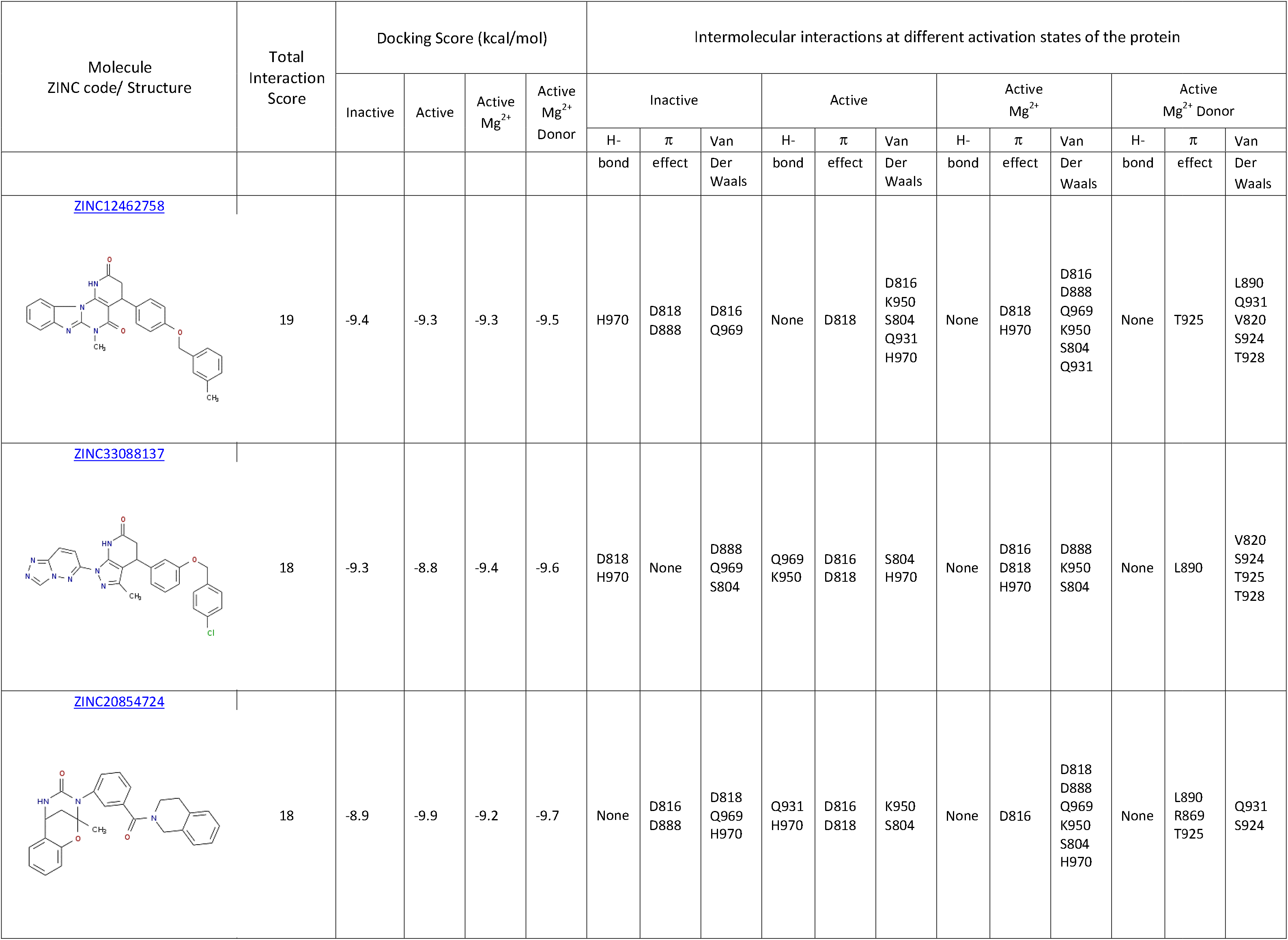
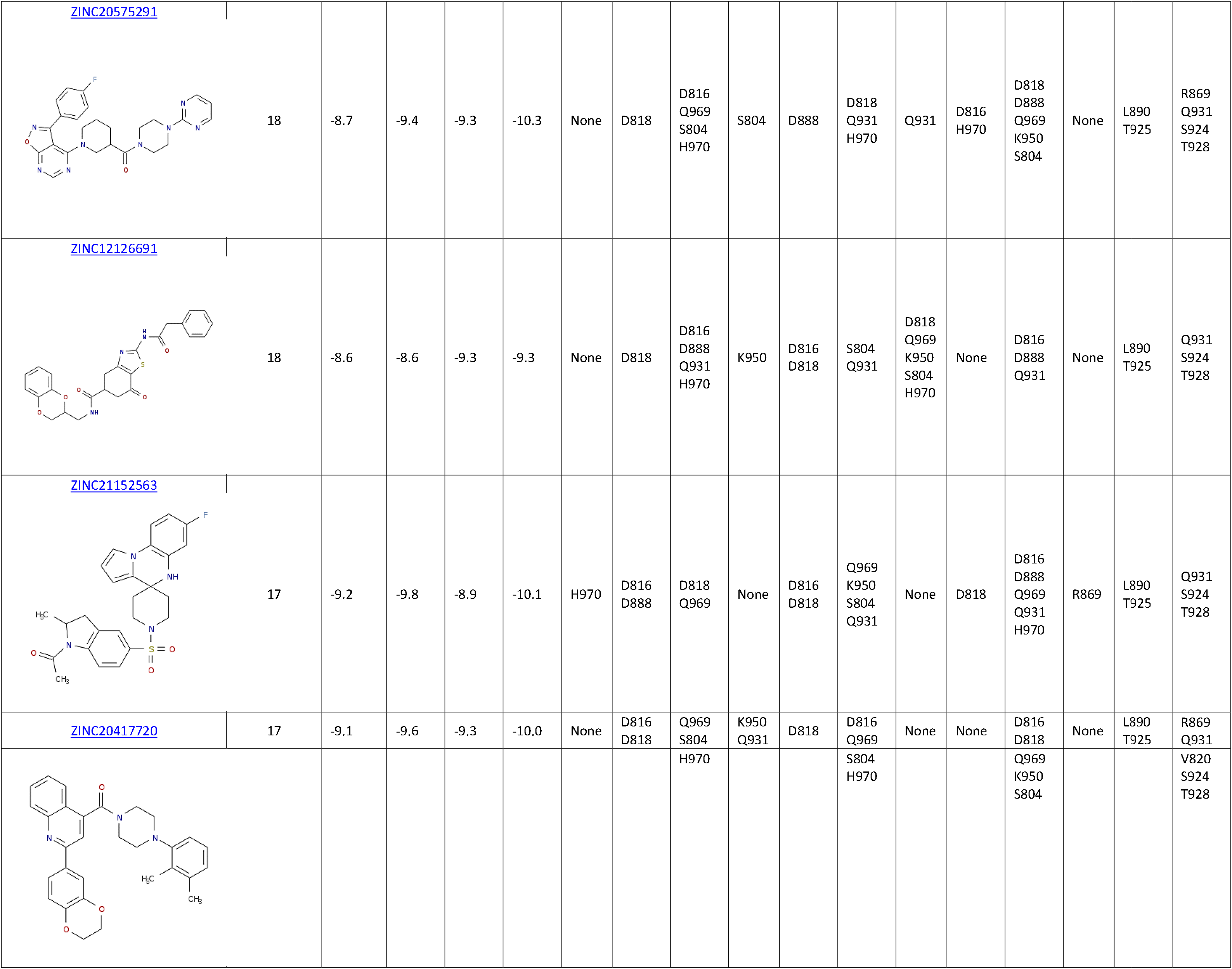
Information associated with potential inhibitors for OAS3. The total interaction score was calculated as the sum of the interaction scores per residue (Figure 2). The intermolecular interactions presented only consider amino acids involved in the binding of the donor or the acceptor substrate. Web pages with details about the candidate molecules and biological tests are attached to each ZINC code. This information was retrieved from [54].

